# Altered neural activity in the mesoaccumbens pathway underlies impaired social reward processing in *Shank3*-deficient rats

**DOI:** 10.1101/2023.12.05.570134

**Authors:** Marie Barbier, Keerthi Thirtamara Rajamani, Shai Netser, Shlomo Wagner, Hala Harony-Nicolas

## Abstract

Social behaviors are crucial for human connection and belonging, often impacted in conditions like Autism Spectrum Disorder (ASD). The mesoaccumbens pathway (VTA and NAc) plays a pivotal role in social behavior and is implicated in ASD. However, the impact of ASD-related mutations on social reward processing remains insufficiently explored. This study focuses on the *Shank3* mutation, associated with a rare genetic condition and linked to ASD, examining its influence on the mesoaccumbens pathway during behavior, using the *Shank3*-deficient rat model. Our findings indicate that *Shank3*-deficient rats exhibit atypical social interactions and have difficulty adjusting behavior based on reward values, associated with modified neuronal activity of VTA dopaminergic and GABAergic neurons and reduced dopamine release in the NAc. Moreover, we demonstrate that manipulating VTA neuronal activity can normalize this behavior, providing insights into the effects of *Shank3* mutations on social reward and behavior, and identify a potential neural pathway for intervention.

## Introduction

Social behaviors play a pivotal role in shaping our lives and fostering a sense of belonging in society. Deficits in these behaviors are a hallmark feature of several psychiatric and neurodevelopmental disorders, including autism spectrum disorder (ASD)^1^. The mesoaccumbens reward pathway is involved in regulating certain aspects of social behavior and a growing body of clinical imaging evidence has pointed towards impaired function of this pathway in ASD^2–9^. Nevertheless, it remains uncertain whether and in what manner ASD genetic risk factors directly impact the assessment of reward value associated with social cues, as only a limited number of studies have delved into this. Given the lack of effective pharmacological interventions for addressing social behavior deficits, there is a pressing need to elucidate the underlying pathophysiological mechanisms.

The mesoaccumbens pathway connects the ventral tegmental area (VTA) and the nucleus accumbens (NAc), two key brain regions involved in reward processing. The VTA is comprised of diverse cell populations that play distinct roles in motivated behaviors and reward processing, with the majority of these cells being dopaminergic (DAergic) (∼60% of VTA cells) and GABAergic (35% of VTA cells) neurons^10–19^. VTA-DA neurons project to several limbic structures including the prefrontal cortex (PFC) and the NAc^13,20–23^. An early neuroimaging study in individuals with ASD showed reduced release of dopamine in the PFC^24^ and significant increase in dopamine transporter binding throughout the brain^25^. Furthermore, imaging studies involving task performance showed that individuals with ASD display a reduction in phasic striatal DA events evoked by social stimuli^8,26^. Together, these findings suggest that functional changes in the dopaminergic system may be implicated in the pathophysiology of ASD.

One notable genetic condition associated with ASD is Phelan-McDermid syndrome (PMS)^27,28^. This rare disorder is caused by mutations or deletions in the *SHANK3* gene, which encodes for a critical scaffolding protein in the core of the postsynaptic density^29–35^. Individuals with PMS present with a wide range of developmental, cognitive, and medical abnormalities^36,37^. Those include generalized developmental delay with intellectual disability of variable severity, absent or delayed speech, hypotonia, motor skill deficits, seizures, gastrointestinal problems, renal malformations, and non-specific dysmorphic features^37–39^. Psychiatric symptoms are prominent in a portion of individuals with PMS, including atypical bipolar disorder, catatonia, attention-deficit hyperactivity disorder^40–43^, and ASD^36,37^. Approximately 63% of individuals with PMS meet DSM-5 criteria for ASD^37^ and up to 2% of individuals with ASD have *SHANK3* haploinsufficiency^44,45^.

To date, there have been no clinical studies examining the involvement of the mesoaccumbens pathway in the PMS phenotype or investigating how social and non-social reward processing may be affected. Despite the lack of clinical studies, the creation and characterization of rodent models with a *Shank3* gene mutation, have consistently demonstrated that *Shank3* deficiency has a detrimental effects on synaptic transmission and plasticity within several brain regions including the striatum and hippocampus; two brain regions that receive DAergic projections from the VTA^28,34,46–52^. Deficits in synaptic transmission were also observed in the VTA of mice following early postnatal down expression of the Shank3 protein within the VTA. Specifically, Bariselli and colleagues, demonstrated that down expression of Shank3 in the VTA impairs the maturation of excitatory synapses onto VTA-DA and VTA-GABA neurons, leading to a decrease in dopamine neuron activity and an increase in GABAergic neural activity in the VTA^47^.

Expanding on these results, in our current study we asked what are the consequences of *Shank3* deficiency on real-time activity of VTA neurons, as well as the release of DA in the NAc during social and non-social behaviors. To address our questions we used a previously validated *Shank3*-deficient rat model^49^, which carries a mutation in the *Shank3* gene that is similar to a human *SHANK3* mutation^53^. This model allowed us to investigates the consequences of *Shank3* deficiency in the entire organism during early developmental stages, a condition that closely resembles the patient’s condition. We utilized fiber photometry tools coupled with calcium or DA sensors to accurately track the activity of DAergic and GABAergic neurons in the VTA, as well as the release of DA in the NAc, in rats during their participation in behavioral tasks. Ultimately, we employed optogenetic techniques to investigate whether altering the activity of VTA neurons could bring about changes in the behavioral phenotype. This allowed us to draw causality between activity of the mesoaccumbens pathway and the observed behavioral phenotype.

## Methods

### Animals

We used adult (8-16 weeks old) male *Shank3*-Heterozygous (*Shank3*-HET), *Shank3*-homozygous/knockout (*Shank3*-KO) and their littermate Wild type (WT) rats. *Shank3*-deficient rats were generated using zinc-finger nucleases on the outbred Sprague–Dawley background, as previously described^49^. Rats were kept under veterinary supervision in a 12 hours reverse light/dark cycle at 22 ± 2°C. Animals were pair-caged with food and water available ad libitum, except during a 48 hours period of food deprivation on the Social vs. Food task when they had access to water but not food. Experiments were conducted during the light phase cycle. All animal procedures were approved by the Institutional Animal Care and Use Committees at the Icahn School of Medicine at Mount Sinai and the University of Haifa.

### Stereotaxic surgeries for viral injections

8-week-old animals were anesthetized with 3-5% isoflurane for induction. Isoflurane was then maintained at 1.5-2.5% with 2% oxygen, using a tabletop vaporizer and a non-breathing circuit. The surgical area was shaved and sterilized before a vertical incision was made along the midline of the skull. After clearing the connective tissue, bregma and lambda were identified, the region of injection was marked, and a small burr hole (50µm) was drilled. The rats were injected unilaterally (for the fiber photometry experiments) or bilaterally (for the optogenetic experiments) with the appropriate viruses. Viruses were loaded into a 10 µl 33G NanoFil syringe (World Precision Instruments, Sarasota, FL, USA) and 0.350 μl of the virus was injected into the VTA (A-P -5.6mm, M-L 0.9mm, D-V 8.0mm) at a rate of 0.1 μl/min. Following injection, the syringe was left in place for 5 min and withdrawn at a rate of 0.2 mm/min. Immediately after, a fiber optic cannula (400um 0.39NA, Cat. CFM14L10, Thor Labs, Newton, New Jersey) was implanted in the same coordinates for the fiber photometry or bilaterally with a 10° angle for the optogenetic experiments. The incision wound was closed using sutures (Ethilon Suture 5-0, Henry Schein, Melville, NY, USA). Animals received intraoperative subcutaneous fluids (Lactated Ringer Solution, Thermo Fisher Scientific, Waltham, MA, USA) and buprenorphine (0.5mg/kg) for analgesia. Additional analgesia was administered subcutaneously every 12 hours for 72 hours post-operatively.

### Viral Vectors

All the viruses used in this study has been previously employed by several other groups^54–60^. To record from VTA-DA neurons, we used a combination of AAV9.rTH.PI.Cre.SV40^56,59^ (Addgene 107788) and AAV9-CAG-FLEX-GCaMP6m.WPRE.SV40^54^ (Addgene 100841). To record from VTA-GABA neurons we used a combination of rAAV-hVGAT1-Cre-WPRE-hGH polyA^60^ (biohippo, Cat no# PT-0346) and AAV9-CAG-FLEX-GCaMP6m^54^ (Addgene 100841). To record from DA release in the NAc, we used a GRAB sensor AAV9-hSyn-DA2m (DA4.4)^58^ (WZ Biosciences). To activate VTA-GABA neurons in WT rats, we used a combination of rAAV-hVGAT1-Cre-WPRE-hGH polyA (biohippo) and AAV9-Ef1a-DIO-hChR2(E123T/T159C)-EYFP^57^ (Addgene, 35509). To activate VTA-DA neurons in *Shank3*-deficient rats, we used a combination of AAV9.rTH.PI.Cre.SV40^56,59^ (Addgene 107788) and AAV9-Ef1a-DIO-hChR2(E123T/T159C)-EYFP^57^ (Addgene 35509). To inhibit VTA-GABA neurons in *Shank3*^-^ deficient rats, we used a combination of rAAV-hVGAT1-Cre-WPRE-hGH polyA^60^ (biohippo) and AAV9-FLEX-Arch-GFP (Addgene 22222)^55^. To inhibit VTA-DA neurons in WT rats, we used a combination of AAV.rTH.PI.Cre.SV40 (AAV9)^56,59^ (Addgene 107788) and AAV9-FLEX-Arch-GFP^55^ (Addgene 22222).

### Behavioral Assays

#### Social/Object vs. Empty Task

Rats were habituated to a testing arena containing two empty compartments on opposite corners of the arena for 15 minutes, as previously described^61^. A social stimulus (novel male Juvenile, 4 weeks old) or a moving object (toy rat) was then placed into one compartment with the opposite compartment left empty (counterbalanced). Compartments had a wire mesh window which allowed for the exchange of visual, auditory, and olfactory information, with very limited physical touch. The subject rat was then allowed to interact for 5 minutes.

#### Social/Object vs. Food Task at satiety and after 48h of food deprivation

Rats were habituated to a testing arena containing two empty compartments on opposite corners of the box for 15 minutes, as previously described^61^. A social stimulus (novel male juvenile, 4 weeks old) or a moving object (toy rat) was then placed into one compartment and food pellets were placed at the opposite compartment (counterbalanced). The subject rat was then allowed to explore the arena for 5 minutes. The task was repeated after 48 h of food deprivation with new juveniles and both compartments on the opposite corners randomly rearranged to avoid habituation and learning.

#### Food consumption Task at 48h of food deprivation

After 48h of food deprivation, rats were habituated to a testing arena containing two empty compartments on opposite corners of the box for 15 minutes. Three tests were carried out:

1. Food pellets were placed in the middle of the open field. The subject rat was allowed to explore and consume the pellets for 5 minutes.
2. A social stimulus (novel male juvenile, 4 weeks old) was placed into one compartment with the opposite compartment left empty, and food pellets were placed in the middle of the open field. The subject rat was then allowed to explore and consume the pellets for 5 minutes.
3. A moving object (toy rat) was placed into one compartment with the opposite compartment left empty, and food pellets were placed in the middle of the open field. The subject rat was then allowed to explore and consume the pellets for 5 minutes.

The amount of grams consumed was determined by measuring the initial weight of the food pellets and then subtracting the weight of the remaining pellets after testing.

### Fiber Photometry Recording and Analysis

Calcium signal recording was conducted using fiber photometry (RZ10x; Tucker-Davis Technologies, FL, USA). The 465 and 405 nm LEDs were driven at 300 mA and 100 mA respectively, with the power at the tip of the fiber optic cannula determined to be at 80-120 uW. A USB camera (C615 portable HD webcam, Logitech®) was placed at the top of the acoustic chamber and connected to a computer for video recording of the subject rat behavior (∼15 frames per second). The video clip recorded by the USB camera was synchronized to the calcium signal as described in the TDT manual (https://www.tdt.com/docs/synapse/hardware/video-processors/#rv2-video-tracker). TDT Synapse software (TDT) was used for recording the signal channel (excitation 470 nm), the isobestic control channel (405 nm) and the digital channel receiving the camera strobes. Three weeks following viral injection and fiber implantation, rat subjects were first habituated to the testing arena for 15 minutes. Calcium signals were then recorded for 5 min before and 5 min after introducing the stimulus/stimuli, by connecting a fiber optic patch cord (400 um, 0.48 NA, Doric lenses, Quebec, Canada) to the fiber optic cannula. Calcium signal data was analyzed following a previously published pipeline, using a custom-written MATLAB Script ^62^. First, we fitted the 405 channels onto 465 channels to detrend signal bleaching and any movement artifacts, according to the manufacturer’s protocol (https://github.com/tjd2002/tjd-shared-code/blob/master/matlab/photometry/FP_normalize.m). Next, the signal was aligned to the video recording using the timestamps recorded by the digital port of the RZ10× system. Calcium signal was aligned to each event and normalized using *z* score (0.1 s bins; zdF/F), where the 2-s pre-event period served as the baseline. The duration of the pre-event period was determined to be 2 seconds since our analysis revealed that the majority of transitions between the two stimuli took 2-3 seconds (**Extended data Figs. 2C, 2F, 6I, and 6J**). Heatmaps were created to depict the fluctuation in fluorescence signals (zdF/F) during the period ranging from 2 seconds before to 5 seconds after each interaction bout. It is important to note that certain rats never engaged with one of the stimuli, resulting in a lack of photometry data. Consequently, these rats were not included in the heatmap, causing variations in the number of rows (representing rats) between the two stimuli. Signal changes were quantified for relevant time intervals as the corresponding areas under the curve for the averaged z-scores, which was calculated using GraphPad software (GraphPad Prism, San Diego, CA, USA) using the ‘area under the curve’ function.

### Optogenetics and behavior

Optogenetic activation or inhibition experiments were conducted three weeks following viral injections and fiber implantations. Rats were first habituated to the testing arena for 15 minutes. Then a splitter fiber optic patch cord (400 um, 0.37 NA, Doric lenses, Quebec, Canada) was connected to the implanted bilateral fibers (CFM14L10, Thorlabs Inc.) through sleeves (F210-3012, Doric lenses) to deliver blue (activation, 465 nm, 5Mw, 5 Hz for the DA neurons with a 20ms pulse width and 10 Hz for the vGAT neurons with a 10ms pulse width) or yellow (inhibition, continuous, 560 nm, 5mW) light (Doric lenses). The stimuli were then introduced, and the behavior was recorded for 5 min. The light was delivered whenever the test rat approached the social stimulus on the Social vs. Empty task at satiety or the Social vs. Food task at 48 hours of food deprivation (light ON). Rats were randomly assigned, with half starting the first round of testing with the lights ON and half with the lights OFF. Then, on an independent day for the second round of testing, those that had the lights ON during the first round switched to having the lights OFF, and vice versa.

### Behavioral analysis

All behaviors were scored and quantified using TrackRodent, an open-source Matlab based automated tracking system (https://github.com/shainetser/TrackRodent) that uses a body-based algorithm (*WhiteRatBodyBased15_7_15* in the TrackRodent interface)^61,63^. The behavioral traces and heat-maps were obtained using a MATLAB custom-made code (https://zenodo.org/records/10222543). Videos were de-identified in order to keep the experimenter blinded to the treatment groups while setting up the analysis. We pooled together all behavioral data for each task, which allowed us to effectively increase the sample size. This was made possible because the different cohorts of rats (each administered with different virus combinations) were all assessed on identical tasks. This enhanced the statistical power of our findings and allowed us to gain a more comprehensive understanding of the behavioral characteristics. However, to maintain clarity and transparency, we present each cohort’s data separately in the Extended Data, and statistical analysis for each cohort is listed in the **Extended Data Table. 1**

#### Investigation time

Behavioral analysis was done after correcting the raw behavioral data by considering any gap of <0.5 s in investigation of a given stimulus as part of the same investigation bout. Investigation time was calculated in 1-minute bin across all tests.

#### Investigation and number of bouts

Given that we have previously reported that longer bouts are associated with social interaction, rather than general exploration^61^, we categorized the different investigation bouts according to their length (<6 sec, 6-19 sec and >19 sec), and calculated investigation time and number of each for each duration category, as done before^61^.

#### Transitions

A transition between stimuli was defined as the time point when investigation of a new stimulus (relative to the other stimulus) started. The mean rate of transitions was calculated at 1-min bins.

#### Interval duration

We evaluated intervals between bouts in which the animal shifted from one stimulus to the other. We counted the number of intervals according to their length and calculated their percentage for each interval duration.

### Histology

After the completion of behavioral testing, rats were deeply anesthetized with 4-5% isoflurane in an induction chamber. Once reaching a surgical plane of anesthesia, rats were perfused transcardially at a rate of 40ml/min with 0.9% NaCl, followed by ice-cold 4% paraformaldehyde (PFA) fixative in 0.1 M phosphate buffer saline (PBS) at pH 7.4. Brains were extracted, post-fixed in 4% PFA for 12 hours at 4°C, and cryoprotected by immersion in a 15% sucrose solution in 0.1 M PBS for 24 hrs at 4°C. Prior to sectioning, brains were flash frozen in isopentane and stored at -20°C. Brains were sectioned with a cryostat (Leica CM 1860 Leica Biosytems, Buffalo Grove, IL, USA) and a series of 30μm thick coronal sections of the VTA or NAc region, collected in a cryoprotective solution (1:1:2 glycerol/ethylene glycol/phosphate buffer saline) and stored at 4°C.

### Immunofluorescent staining

Brain sections were rinsed in 1X PBS with 0.05% Triton X-100 (PBS-T), then incubated in a primary antibody anti-tyrosine hydroxylase (1:2000, mouse, monoclonal, MAB318, EMD Millipore®) and/or anti-GFP (1:2000, chicken, A10262, invitrogen®) overnight at 4°C in 10% nonfat dry milk and 1% bovine serum albumin (BSA) in 0.03% Triton X-100 in 1xPBS. The following day, sections were washed with 1X PBS with 0.05% Triton X-100 (PBS-T) and incubated in secondary antibody (Cy3, Donkey anti-mouse, code: 711-165-152, and Alexa Fluor® 488, Donkey anti-chicken, code:703-545-155, Jackson Immuno Research®) (1:1000 in 0.5% Triton X-100 in PBS) for 2 hours at room temperature. Sections were then rinsed and mounted with VECTASHIELD Antifade Mounting Medium with DAPI (cat. no. H-1200, Vector Labs, Burlingame, CA, USA).

### RNAscope

Rat *vgat1 (slc32a1)* probe was purchased from ACDBio. After perfusion, extraction, post-fixation and sucrose immersion, brains were flash frozen in a slurry of isopentane and dry ice. Tissue was sectioned at 20µm, mounted on glass slides (SuperFrost Plus Microscope Slides, Fisher Scientific, USA) and frozen at -80°C until the day of experiment. RNAscope was performed following the manufacturer’s protocol (RNAscope Multiplex Fluorescent Reagent Kit, ACDBio, USA). Briefly, tissue sections were thawed at RT for 10min, fixed with 4% PFA for 15min at 4°C, and dehydrated with varying ethanol concentrations. They were then incubated in H_2_O_2_ for 10min and the *vgat1* probe added and incubated for 2 hours in a 40°C oven (HybEZ II Hybridization System, ACDBio, USA). This was followed by an amplification step and then incubation with opal dye (Akoya Biosciences, USA) 570 to visualize the RNA transcripts. Immunohistochemical staining for GFP was performed following the RNAscope experiment with a blocking step (1h, RT) with donkey serum and incubation of tissue with anti-GFP (1:1000, chicken, A10262, invitrogen®) overnight at 4°C followed by incubation with Donkey-anti-Chicken 488 (2h, RT).

### Image acquisition and processing

Immunofluorescent staining and fluorescent in situ hybridization (RNAscope) were imaged on a Zeiss AxioImager Z2M with ApoTome.2 at 10x magnification at the Microscopy and Advanced Bioimaging CoRE at the Icahn School of Medicine at Mount Sinai.

### Statistical analysis

All statistical analyses were performed using GraphPad prism 9.0 software (GraphPad Prism, San Diego, CA, USA). Detailed information about statistical tests is provided in each figure legend and in the Extended Data, Table 1. Image graphics were created using BioRender.com and Adobe Illustrator.

## Results

### *Shank3*-deficient rats exhibit an atypical social interaction that is associated with impaired VTA-DA neural activity

To examine the effects of *Shank3* mutation on social behavior and the associated neural activity within the VTA, we first conducted a comprehensive assessment of the *Shank3*-deficient rats’ behavior using the Social vs. Empty task. In this task, subject rats were presented with two compartments, one that contained a social stimulus (juvenile rat) and the other that was kept empty (**Fig. 1A**). We found that WT rats and their *Shank3*-Het and *Shank3*-KO littermates all spent more time investigating the compartment containing a same-sex juvenile rat stimulus, compared to the empty compartment (**Fig. 1B** and **Extended Data Fig. 1A**). At first glance, this outcome may be misconstrued to suggest that *Shank3*-deficient rats exhibit social behavior that is similar to their WT littermates. However, *Shank3*-KO rats exhibited distinctly elevated time interacting with the social stimuli, when compared to their WT littermates (**Fig. 1B**). They also exhibited significantly higher engagement in long-bouts of social investigation (>19 sec), previously shown to be more associated with social interaction^61^ (**Extended Data Fig. 1B**, upper right panel), and significantly lower engagement with the empty compartment, regardless of the bout length (**Extended Data Fig. 1B**, lower panels), with no change in the total number of bouts (**Extended Data Fig. 1C**). A detailed analysis of the overall interaction characteristics during the 5-minute testing period further re-affirmed that *Shank3*-deficient rats exhibit an atypical social behavior. WT rats showed an increase in transitioning between the stimuli (**Extended Data Fig. 1D,** upper panels), while *Shank3*-HET and KO rats showed no change in the transition rate over the 5-minutes period of testing (**Extended Data Fig. 1D,** middle and lower panels, respectively). This atypical behavior was accompanied with a distinct pattern of investigation across the 5 minutes of testing. Specifically, WT rats exhibited a substantial decline in their engagement with the social stimuli at the second minute of testing, to the extent that they displayed no preference between the social and empty compartments at the third minute (**Fig. 1C and 1D**, left panels). *Shank3*-Het and *Shank3*-KO rats, however, continued to interact with the social stimulus and exhibited almost no interest in the empty compartment throughout the entire 5 minutes of testing (**Fig. 1C and Fig. 1D**, middle and right, respectively). These differences in stimuli preference between genotypes were evident through the statistically significant variations in the preference score during the second and third minutes of testing (**Fig. 1E**). Critically, in a comparable paradigm where we introduced an object (a moving rat toy) instead of a social stimulus (**Fig. 1F**, Object vs. Empty), both WT and *Shank3*-KO rats exhibited a preference for the novel object (**Fig. 1G** and **Extended Data Fig. 1E**), albeit to a lesser degree compared to their preference to the social stimulus in the Social vs. Empty task. While rats from all three genotypes showed a significantly strong preference to the object stimulus during the 1^st^ minute of testing, WT and *Shank3*-Het rats quickly lost interest in investigating the object following the first minute (**Fig. 1H** and **1I**, left and middle panels, respectively), while *Shank3*-KO rats continued to spend more time at the object compartment (**Fig. 1H** and **1I**, right panels and **Fig. 1J**). We observed no significant differences in the cumulative bout number (**Extended Data Fig. 1F**), bout length (**Extended Data Fig. 1G**), or transition pattern (**Extended Data Fig. 1H**) on the Object vs. Empty task. Together, these findings suggest that *Shank3*-KO rats demonstrate an atypical increase in interaction when faced with a single stimulus choice, regardless of whether it involved social or non-social stimuli.

**Figure 1.**
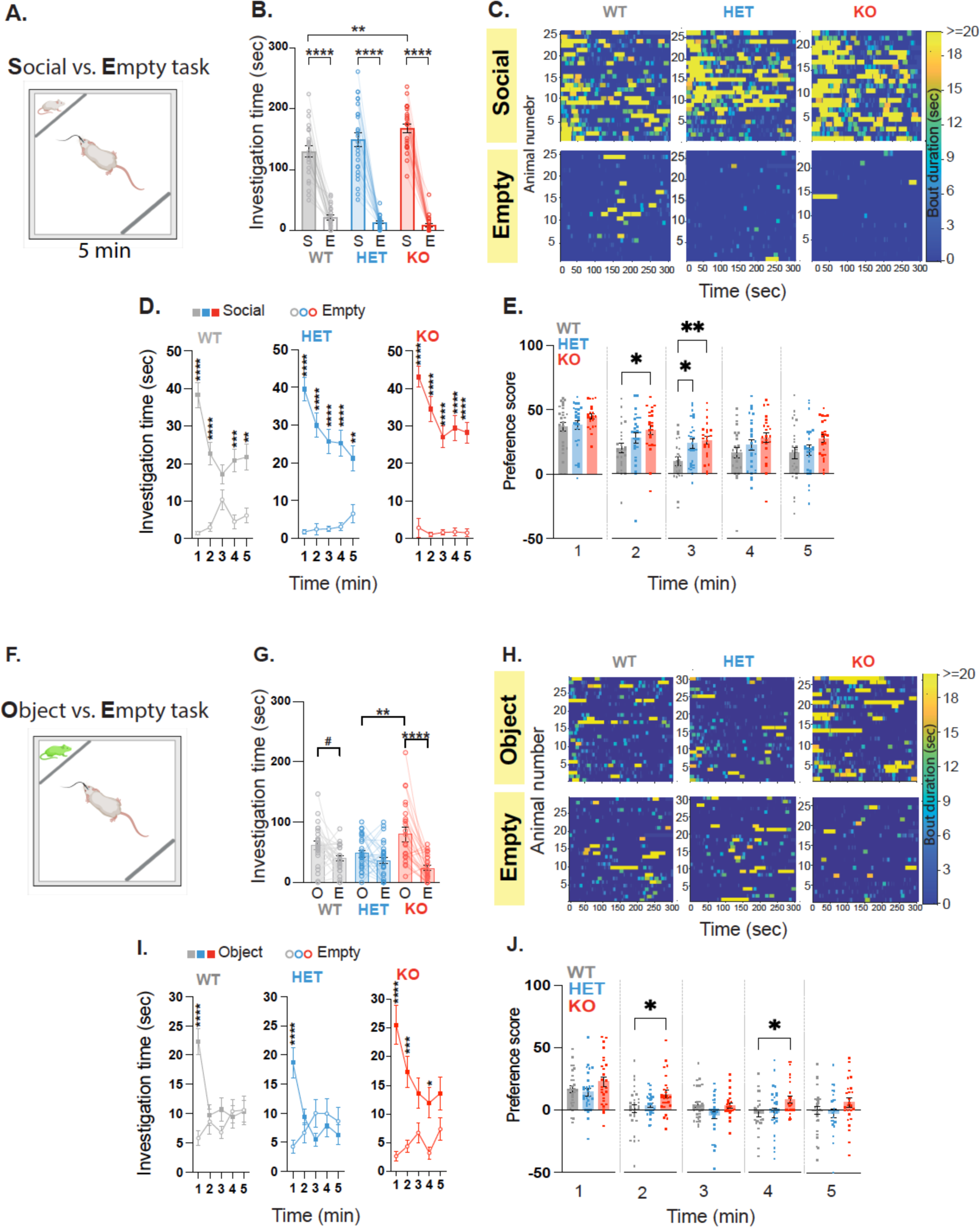
*Shank3*-deficient rats show an atypical behavior when presented with a social or object stimuli. **A**. Behavioral outline for the Social vs. Empty task. Rats encounter either a confined novel same-sex juvenile or an empty compartment. **B.** Mean total investigation time of the social stimulus or the empty compartment. **C.** Heat maps depicting the investigation time of individual rats during the 5-minute test period. Each row represents one rat. **D.** Mean total investigation time for the social stimulus and the empty compartment averaged in 1-minute intervals over the 5-minute test period. **E.** Differences in mean investigation time between the social stimulus and the empty compartment calculated for each minute. B-E: WT, n=25; HET, n=26; KO, n=23. **F**. Behavioral outline for the Object vs. Empty task. Rats encounter either a confined, moving toy-rat or an empty compartment. **G-J.** Similar to B-E, but show behavior during the Object vs. Empty task. G-J: WT, n=29; HET, n=30; KO, n=27. *\*\*\*\*p*<.0001, *\*\*\*p*<.001, *\*\*p*<.01, *\*p*<.05, post hoc tests following the main effect. All error bars represent SEM. Detailed statistical data are provided as a Data file in Extended Data, Table 1.

To record neural activity of VTA neurons during behavior, we employed an *in vivo* fiber photometry calcium imaging approach in conjunction with a genetically encoded calcium indicator (GCaMP). This technique provides a highly sensitive tool to record cell-type specific neuronal activity in behaving animals within deep brain structures^64,65^, such as the VTA^66^. To specifically target VTA-DA neurons, we used a dual viral approach where one virus expressed a Cre-dependent GCaMP6 (AAV9-CAG-FLEX-GCaMP6m) and a second virus expressed Cre recombinase under the control of the tyrosine hydroxylase (TH) promoter, a common promoter used to target DAergic neurons^67^ (AAV9.rTH.PI.Cre.SV40) (**Fig. 2A** and **2B**). Three weeks following viral expression, we recorded GCaMP6 fluorescent signals during the Social vs. Empty task (**Fig. 2C**). We found a significant increase of calcium signals in WT rats during social interaction (**Fig. 2D-F**), reflecting an increase in VTA-DA neural activity that is consistent with previous reports in mice^46,47,66,68^. In contrast, despite displaying seemingly active and even elevated interaction with the social stimuli (**Extended Data Fig. 2A-B**), *Shank3*-Het and KO rats did not exhibit an increase in VTA-DA neural activity during social interaction (**Fig. 2D-F**), and showed a comparable pattern of activity when approaching the object on the Object vs. Empty task (**Fig. 2G-J** and **Extended Data Fig. 2D and 2E**). These results raise the possibility that *Shank3*-deficient rats do not appropriately process the rewarding value of social interactions, due to the lack of VTA-DA neuronal activity, hence are not getting satisfaction out of it.

**Figure 2.**
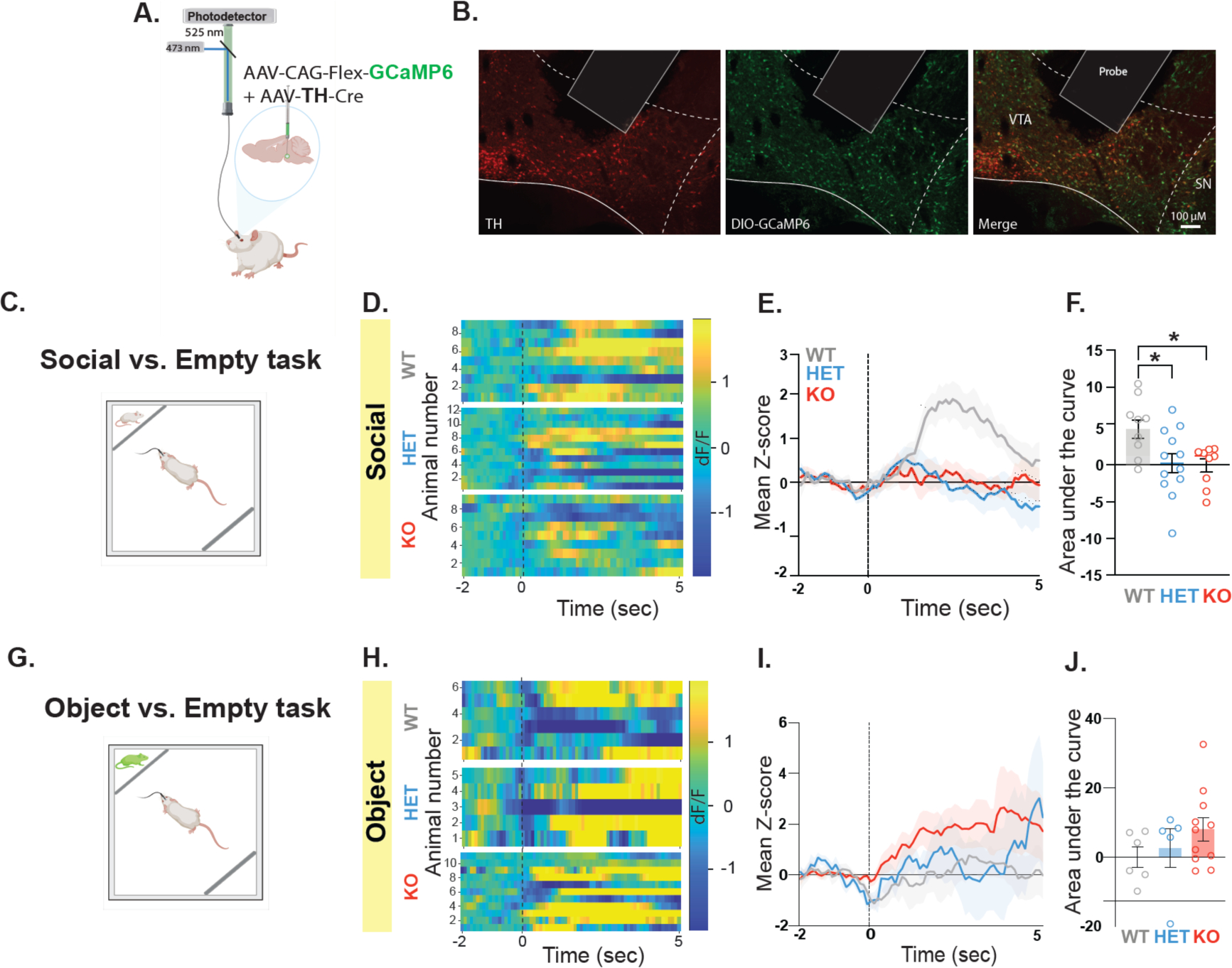
*Shank3*-deficient rats show impaired VTA-DA neural activity during interaction with a social but not an object stimulus. **A.** A schematic of viral injection and fiber photometry setup in the VTA. **B.** Immunohistochemistry staining of a sample VTA section showing TH expression (left panel, red), GCaMP6 (viral) expression (middle panel, green), and overlap (right panel), along with fiber placement (10X magnification; scale bar = 100 μm). **C.** Behavioral outline for the Social vs. Empty task. Rats encounter either a confined novel same-sex juvenile or an empty compartment. **D-F.** GCaMP6 Fiber photometry recordings of VTA-DA neurons during the Social vs. Empty task. WT, n=10; HET, n=12; KO, n=9. **D.** Heat maps illustrate the change in fluorescence signals (dF/F) from 2 seconds before to 5 seconds after each social interaction bout. Each row corresponds to one animal and comprises an average of signals from all interaction bouts during the 5-minute testing period. **E.** Average standardized VTA-DA photometry responses aligned to investigation onset of the social stimulus **F.** The area under the curve, calculated from average standardized traces in (E). **G-J.** Similar to C-F, but show behavioral data and fiber photometry recording during the Object vs. Empty task. WT, n=;11 HET, n=13; KO, n=11. All error bars represent SEM. Detailed statistical analysis is provided as a Data file in Extended Data, Table 1.

### *Shank3*-deficient rats show deficits in shifting their behavior based on the value of the social and non-social rewards

To investigate if *Shank3*-deficient rats can process the value of a rewarding stimuli and alter their behavior based on the change in reward value of the presented stimuli, we used the Social vs. Food task (**Fig. 3A**). This task models a more naturalistic settings where animals are faced with more than one rewarding choice^61,69^. Specifically, rats are presented concurrently with a social (same-sex juvenile rat) and non-social rewarding stimuli (food), each at opposing compartment while at satiety and then tested again on the same paradigm following food deprivation. As social interaction is typically more rewarding than food in caloric satiation state, rats are expected to spend more time investigating the social stimulus than food. However, when food-deprived, the value of food increases, as a result rats are expected to shift their behavior accordingly^61,69^. At satiety, both *Shank3*-deficient rats and their WT littermates exhibited a greater inclination towards engaging with the social stimulus rather than food (**Fig. 3B-E** and **Extended Data Fig. 3A,** left panel) with *Shank*3-HET rats showing a slightly increased interest in the social stimulus (**Fig. 3B**). Also, number of bouts (**Extended Data Fig. 3B**) and total investigation time across bouts (**Extended Data Fig. 3C**) were comparable between genotypes. While *Shank3*-deficient rats displayed no noticeable change in transitions, compared to WT littermates that demonstrated fewer transitions over time (**Extended Data Fig. 3D**), overall we observed no major differences in behavior between genotypes on this task. However, when tested again on the same Social vs. Food task after 48 hours of food deprivation, WT rats shifted their behavior to spend an equal time investigating both stimuli, while *Shank3*-KO rats continued to spend significantly more time investigating the social stimulus (**Fig. 3F** and **Extended Data Fig. 3A,** right panel). Compared to their WT littermates, *Shank3*-KO rats also displayed a significantly higher number and cumulative bout length towards the social stimulus, and a lower number of bouts and cumulative bout length towards the food (**Extended Data Fig. 3E and 3F**) with no substantial difference in the transition pattern between the two stimuli (**Extended Data Fig. 3G**). The pattern of increased investigation exhibited by the *Shank3*-deficient rats was clearly sustained across the five minutes of testing (**Fig. 3G-I)**. To rule out the possibility that the diminished interest of *Shank3*-deficient rats in interacting with food is due to reduced appetite, we subjected the rats to a 48-hour food deprivation period. Afterward, we gave them unrestricted access to food without any additional stimuli in the arena or while presenting them with either a novel juvenile or a moving toy object. *Shank3*-deficient rats consumed food at normal levels when no stimuli were introduced into the testing arena and when a moving object was present (**Extended Data Fig. 4A** and **4B,** respectively). However, when presented with a social stimulus, *Shank3*-deficient rats displayed a significant decrease in food consumption (**Extended Data Fig. 4C)**. Together, these findings suggest that only when presented with a social stimulus, *Shank3*-deficient rats show impairment in adjusting their behavior based on the value of the presented stimuli.

**Figure 3.**
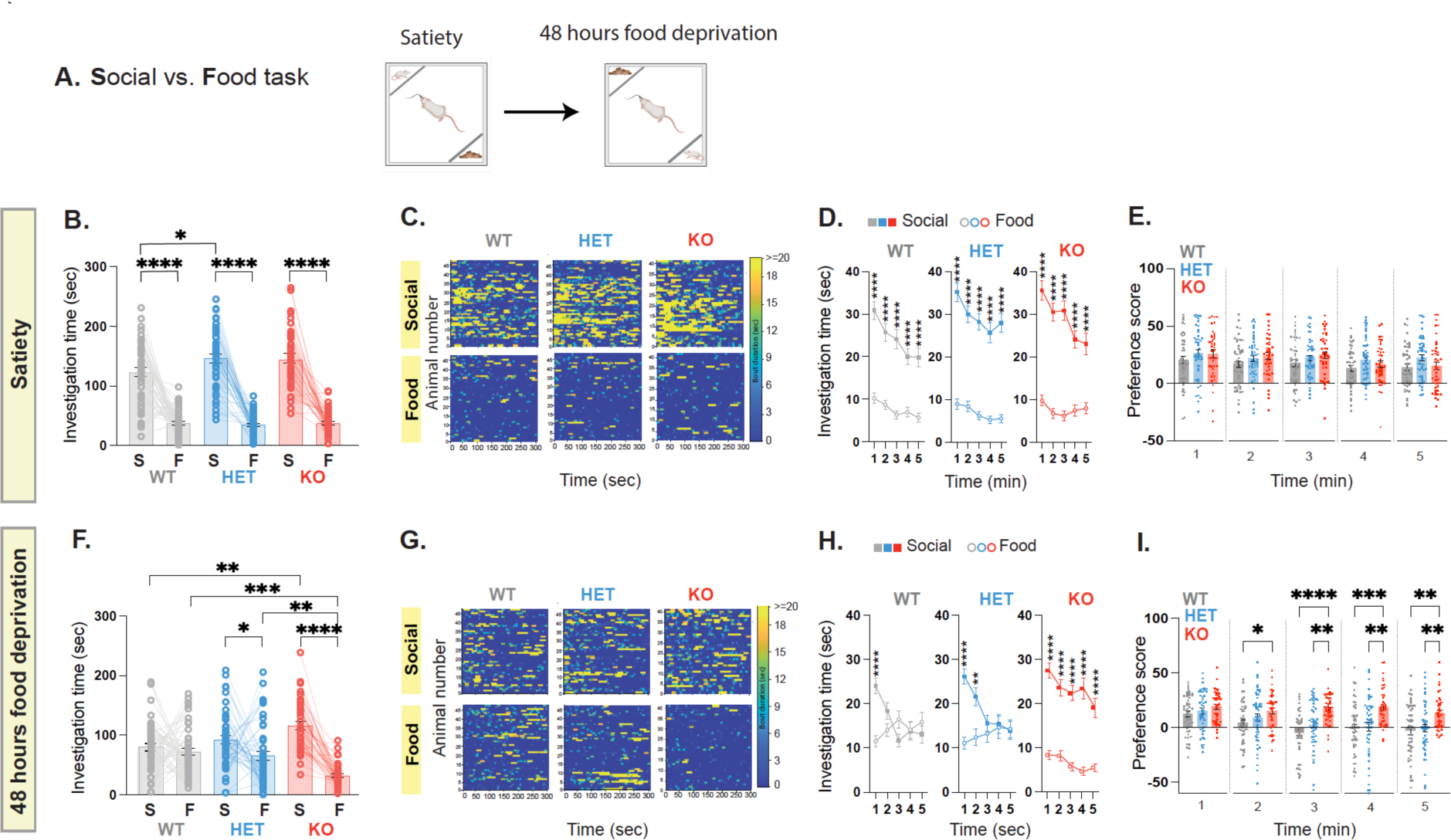
*Shank3*-deficient rats show deficits in shifting their behavior based on the value of the social and non-social rewards. **A.** Behavioral outline for the Social vs. Food task at satiety and after 48 hours of food deprivation. Rats encounter either a confined novel same-sex juvenile or a food-containing compartment. **B.** Mean total investigation time of the social stimulus or the food-containing compartment at satiety. **C.** Heat maps depicting the investigation time of individual rats during the 5-minute test period. Each row represents one rat. **D.** Mean total investigation time for the social stimulus and the food-containing compartment averaged in 1-minute intervals over the 5-minute test period. **E.** Differences in mean investigation time between the social stimulus and the food-containing compartment for each minute**. F-I.** Similar to B-E, but show behavior after 48h of food deprivation. WT, n=47; HET, n=47; KO, n=41. *\*\*\*\*p*<.0001, *\*\*\*p*<.001, *\*\*p*<.01, *\*p*<.05, post hoc tests following the main effect. All error bars represent SEM. Detailed statistical data are provided as a Data file in Extended Data, Table 1.

To further substantiate our findings, we investigated the behavior of the rats using the Object vs. Food task. In this task, rats were simultaneously exposed to both an object and food when they were satiated (**Fig. 4A**). This setup ensured that the object and food were of equal or no reward value. *Shank3*-deficient rats and their WT littermates showed no preference to either the object or food compartments at satiety (**Figs. 4B-E** and **Extended Data Fig. 5A**, left panel). Furthermore, there were no substantial difference across genotypes in the number of bouts (**Extended Data Fig. 5B**), in the cumulative bout investigation time (**Extended Data Fig. 5C**), or in the transition pattern (**Extended Data Fig. 5D**). Importantly, similar to their WT littermates, *Shank3*-*deficient* rats continued to exhibit a typical behavior also following food deprivation, showing preference to food over object (**Fig. 4F-I**), as well as similar number of bouts and cumulative bout investigation time (**Extended Data Fig. 5E** and **Extended Data Fig. 5F,** respectively), and transition patterns (**Extended Data Fig. 5G**). Taken together, our findings suggest that the atypical behavior observed in the *Shank3*-deficient rats pertain specifically to their ability to adjust their behavior based on the rewarding values of two competing stimuli. These impairments are context-specific, becoming apparent when confronted with social stimulus alongside another stimulus of competing value.

**Figure 4.**
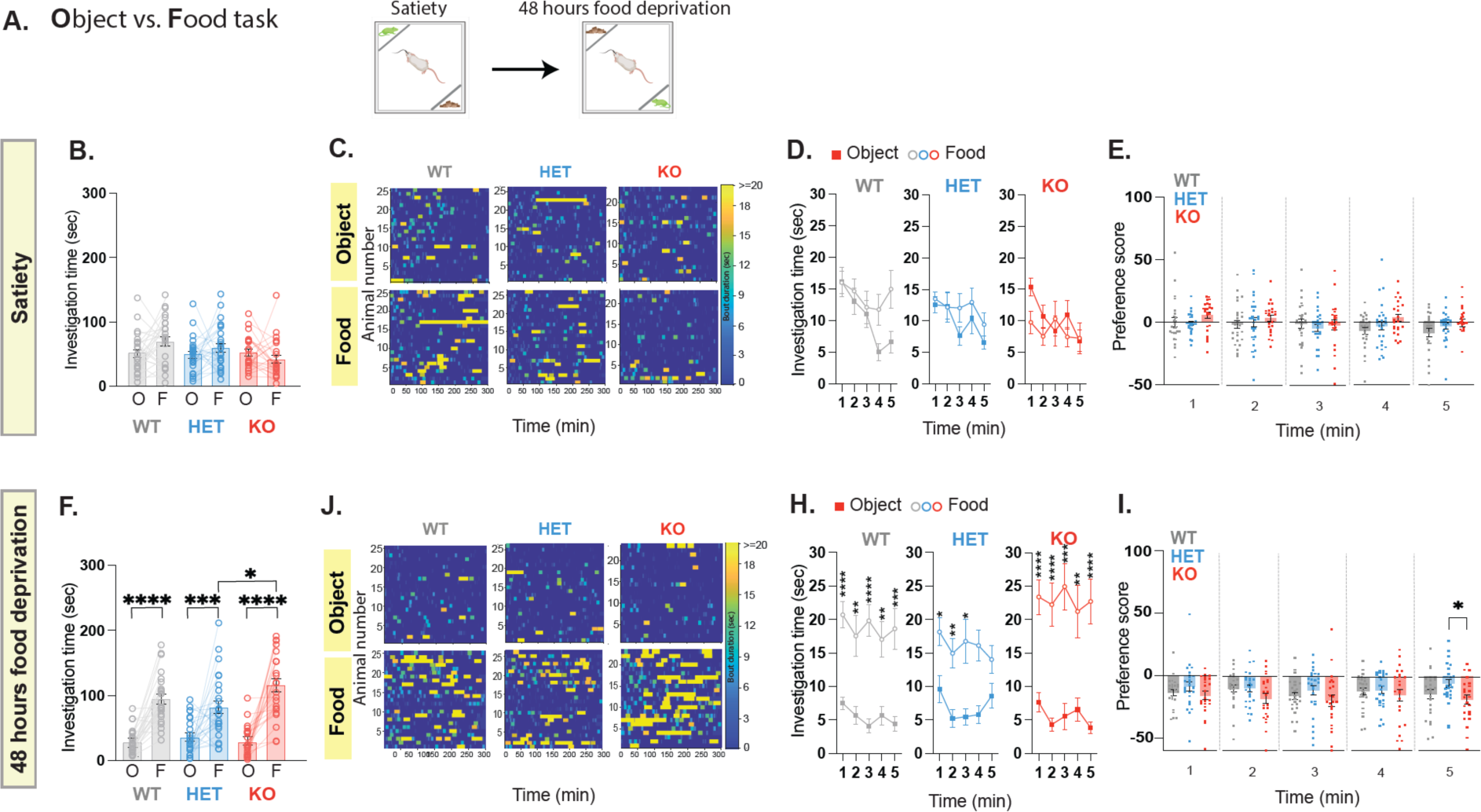
*Shank3-*deficient rats exhibit a typical behavior when presented with an object stimulus. **A.** Behavioral outline for the Object vs. Food task at satiety and after 48 hours of food deprivation. Rats encounter either a confined moving rat toy or a food-containing compartment. **B.** Mean total investigation time of the object stimulus or the food-containing compartment at satiety. **C.** Heat maps depicting the investigation time of individual rats during the 5-minute test period. Each row represents one rat. **D.** Mean total investigation time for the object stimulus and the food-containing compartment averaged in 1-minute intervals over the 5-minute test period. **E.** Differences in mean investigation time between the object stimulus and the food-containing compartment for each minute**. F-I.** Similar to B-E, but show behavior after 48h of food deprivation. WT, n=25; HET, n=26; KO, n=23. *\*\*\*\*p*<.0001, *\*\*\*p*<.001, *\*\*p*<.01, *\*p*<.05, post hoc tests following the main effect. All error bars represent SEM. Detailed statistical data are provided as a Data file in Extended Data, Table 1.

### *Shank3*-deficient rats show impairment in VTA-DA neural activity during social but not food or object interaction

To examine if the observed atypical behavior is correlated with changes in VTA-DA neural activity, we recorded neural activity during both the Social vs. Food (**Fig. 5A**) and Object vs. Food tasks (**Fig. 5H**) during the satiety state, using GCaMP6 signals for recording, while simultaneously confirming that the previously observed behavioral phenotype is replicated in this cohort of rats (**Extended Data Fig. 6A-H**). Our analysis revealed that VTA-DA neural activity increased during social interaction in WT rats, whereas no such increase was observed in *Shank3*-Het or *Shank3*-KO rats (**Fig. 5B-D**). Notably, this lack of response was observed only during interaction with a social stimulus, as VTA-DA neural activity was elevated in all three genotypes when rats explored the food compartment (**Fig. 5E-G**). Furthermore, when recording during the Object vs. Food task, we observed no statistically significant differences across genotypes, regardless of the stimulus (**Fig. 5I-N**). These results reaffirm our earlier findings that, although *Shank3*-deficient rats spend more time engaged in social interaction, their distinctive social interaction characteristics coupled with the absence of an increase in VTA-DA neural activity, suggest ineffective processing of the rewarding aspect of social interaction. Furthermore, these findings emphasize that the lack of enhanced VTA-DA activity in behaving rats is specifically notable during social interaction.

**Figure 5.**
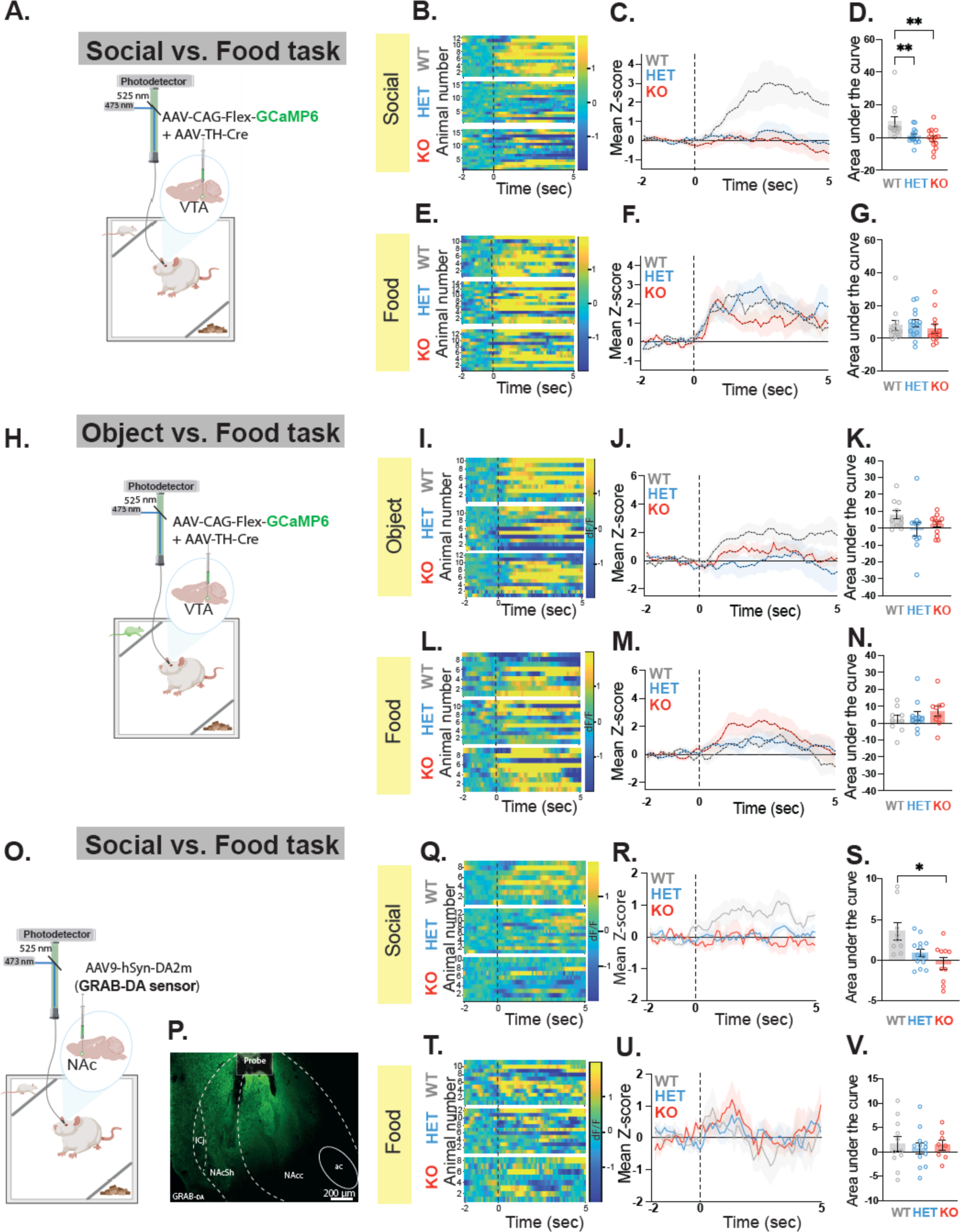
*Shank3*-deficient rats show impaired VTA-DA neural activity during interaction with a social but not an object or food stimulus. **A.** A schematic of viral injection and fiber photometry setup in the VTA. **B-G.** GCaMP6 fiber photometry recordings of VTA-DA neurons during the Social vs. Food task. **B.** Heat maps illustrate the change in fluorescence signals (dF/F) from 2 seconds before to 5 seconds after each interaction bout for the social stimulus. Each row corresponds to one animal and comprises an average of signals from all interaction bouts during the 5-minute testing period. **C.** Average standardized VTA-DA photometry responses aligned to investigation onset of the social stimulus. **D.** The area under the curve, calculated from average standardized traces in (C). **E-G.** Similar to B-D, but show fiber photometry recording when the rats explore the food. WT, n=14; HET, n=14; KO, n=16. **H-N.** Similar to A-G, but shows behavioral data and fiber photometry recording during the Object vs. Food task. WT, n=10; HET, n=12; KO, n=13. **O.** A schematic of viral injection and fiber photometry setup in the NAc. **P.** Immunohistochemistry staining of a sample NAc section showing GRAB^DA^ sensor expression (in green, 10X magnification; scale bar = 200 μm). **Q-V.** Similar to B-G, but show fiber photometry recording of GRAB^DA^ sensor in the NAc during the Social vs. Food task. WT, n=10; HET, n=14; KO, n=10. *\*\*p*<.01, *\*p*<.05, post hoc tests following the main effect. All error bars represent SEM. Detailed statistical data are provided as a Data file in Extended Data, Table 1.

### *Shank3*-deficient rats show impaired DA release in the NAc during social interaction

To investigate whether the impaired behavior and the VTA-DA neural activity deficits are also associated with disrupted DA release in the NAc during social interaction, we utilized the G protein-coupled receptor [GPCR]-activation-based-DA (GRABDA) sensor^70^ in combination with fiber photometry to examine changes in NAc-DA levels during social interaction in an independent cohort of rats (**Extended Data Fig. 7A-D**). Specifically, we injected the AAV9-hSyn-DA2m (DA4.4) into the NAc and implanted an optic fiber just above the injection site (**Fig. 5O** and **5P)**. Three weeks following viral expression, we recorded the fluorescent signals of GRAB_DA_ during the Social vs. Food task when animals were satiated. Our findings showed that in WT rats, social interaction was accompanied by elevated DA release in the NAc, whereas this effect was not observed in the *Shank3*-*deficient* rats (**Fig. 5Q-S**). Less increase was associated with exploration of the food compartments in all genotypes (**Fig. 5T-V**). Together, these results suggest that the reduced neural activity in the VTA-DA neurons of *Shank3*-deficinet rats aligns with a decrease in DA release in downstream VTA regions, notably the NAc, specifically during social interaction.

### *Shank3*-*deficient* rats show abnormal increase in VTA-GABAergic neural activity during social interaction

To investigate the impact of *Shank3* mutation on neural activity of VTA-GABA neurons during social interaction, we injected into the VTA the AAV9-CAG-FLEX-GCaMP6m virus, which was responsible for expressing GCaMP6 in a Cre-dependent manner and the rAAV-hVGAT1-Cre-WPRE-hGH polyA virus, which expressed the Cre recombinase under the control of the vesicular GABA transporter (vGAT) promoter, commonly utilized to target GABAergic neurons^60^ (**Fig. 6A** and **6B**). Our behavioral findings from this independent cohort of rats replicated those from other cohorts (**Extended Data Fig. 8A** and **8B)**. Our recording revealed that while WT rats exhibited a slight increase in VTA-GABA activity when approaching the social stimulus, both *Shank3*-Het and *Shank3*-KO rats displayed a significantly higher increase in VTA-GABA activity (**Fig. 6C-E**). During the Social vs. Food task (**Fig. 6F** and **Extended Data Fig. 8C-F**), we observed a comparable trend of increased VTA-GABA activity, noticeable only when the rats approached the social, but not the food stimulus (**Fig. 6 G-I** and **Fig. 6J-L**, respectively). Finally, we found no differences in VTA-GABA activity patterns between genotypes in either the Object vs. Empty task (**Extended Data Fig. 9A-F**) or Object vs. Food task (**Extended Data Fig. 10A-K**). Taken together, these findings demonstrate that *Shank3*-deficient rats exhibit an atypical increase in VTA-GABA neural activity that is specific to social interaction and not generalized across all stimuli.

**Figure 6.**
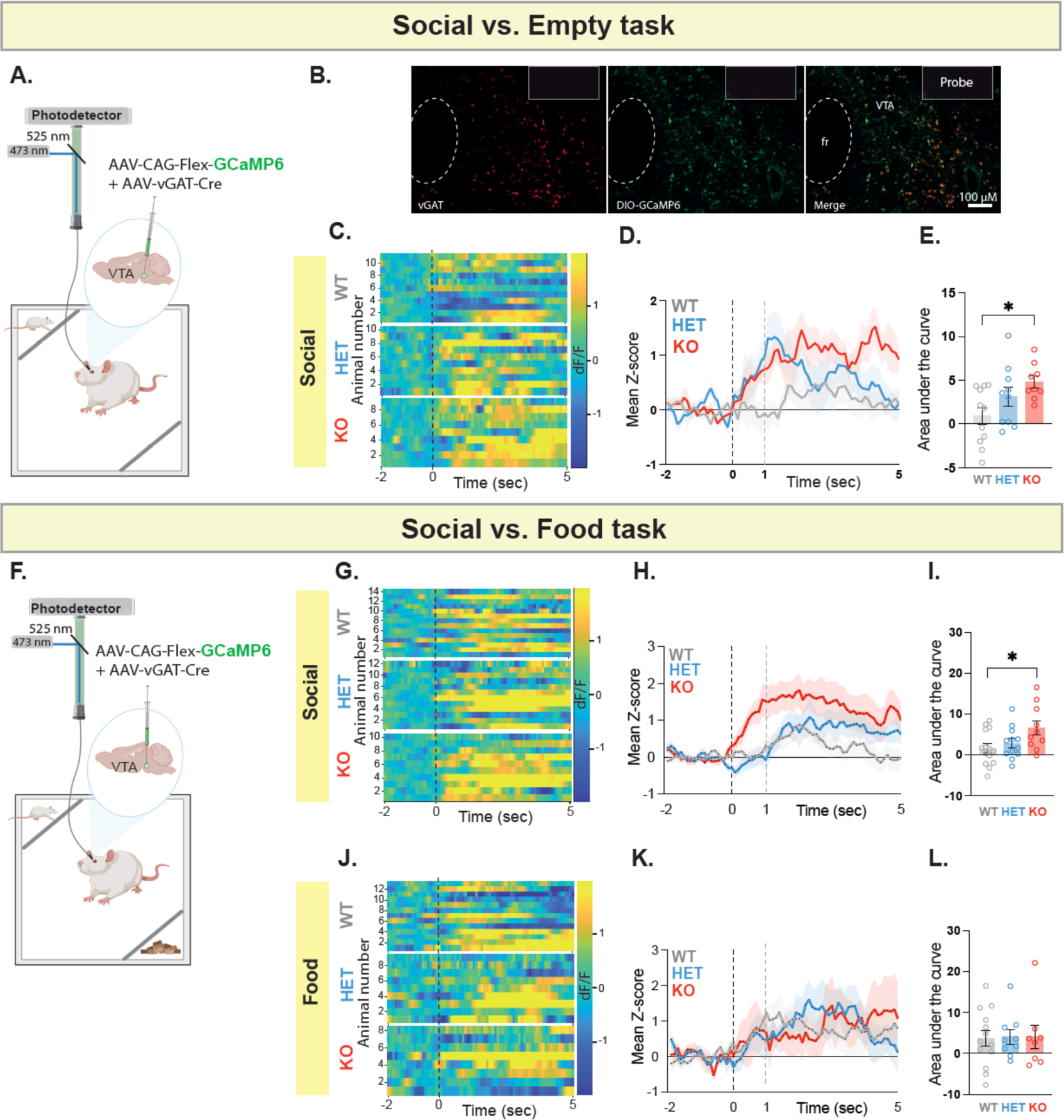
*Shank3*-deficient rats show abnormal increase in VTA-GABA neural activity during social interaction. **A.** A schematic of viral injection and fiber photometry setup in the VTA. **B.** Immunohistochemistry staining of a sample VTA section showing vGAT expression (left panel, red), GCaMP6 (viral) expression (middle panel, green), and overlap (right panel), along with fiber placement (10X magnification; scale bar = 100 μm). **C.** Heat maps illustrate the change in GCaMP6 fluorescence signals (dF/F) from 2 seconds before to 5 seconds after each interaction bout for the social stimulus. Each row corresponds to one animal and comprises an average of signals from all interaction bouts during the 5-minute testing period. **D.** Average standardized VTA-GABA photometry responses aligned to investigation onset of the social stimulus. **E.** The area under the curve, calculated from average standardized traces in (D). WT, n=11; HET, n=10; KO, n=9. **F-L.** Similar to A-E, but show fiber photometry recording during the Social vs. Food task at Satiety. WT, n=14; HET, n=12; KO, n=10. *\*p*<.05. All error bars represent SEM. Detailed statistical analysis is provided as a Data file in Extended Data, Table 1.

### Optogenetic activation of VTA-DA neurons during social interaction normalizes the atypical behavior in *Shank3*-KO rats

In light of our findings, we next tested whether optogenetic activation of VTA-DA neurons could normalize the atypical behavioral phenotype in *Shank3*-KO rats to a level comparable to that of their WT littermates. To selectively activate VTA-DA neurons, we injected the AAV-TH-Cre virus along with the AAV9-Ef1a-DIO-hChR2(E123T/T159C)-EYFP, which expresses a Cre-dependent Channelrhodopsin (ChR2), into the VTA (**Fig. 7A**). In parallel, we carried out a complementary experiment on WT littermate rats to investigate whether inhibition of VTA-DA neurons could elicit an atypical behavioral phenotype similar to that we observed in *Shank3*-deficient rats. For this purpose, we injected the AAV-TH-Cre virus along with the AAV9-FLEX-Arch-GFP virus into the VTA of WT rats (**Fig. 7B**). We found that activating VTA-DA neurons (light-ON) in *Shank3*-KO rats during social interaction on the Social vs. Empty task resulted in a reduction in investigation time of the social compared to the empty compartment and an overall decrease in investigation of the social stimulus (**Fig. 7C-E**). When VTA-DA neurons were inhibited in WT littermate rats (light-ON), there was no significant effect on behavior when examining the total investigation time of the rats (**Fig. 7F**). Nevertheless, it influenced the pattern in which the rats investigated and expressed preference between the social and empty compartments (**Fig. 7G** and **7H**), particularly during the first minute, where we noted a trend in the significance of the preference score (**Fig. 7H**). When tested on the Social vs. Food task, specifically 48 hours after food deprivation, where *Shank3*-KO rats still showed strong preference to the social stimulus despite food deprivation (**Fig. 7I** and **7J**, light-OFF), we found that activating VTA-DA neurons in *Shank3*-KO rats during social interaction increased their interest in the food over the social stimulus (**Fig. 7I** and **7J**, light-ON), which was most notable during the third and fourth minutes on interaction (**Fig. 7K**). Remarkably, in WT rats, where there was a clear preference for food following food deprivation (**Fig. 7L** and **7M**, light-OFF), we found that inhibition of VTA-DA neurons altered that preference (**Fig. 7L** and **7M**, light-ON, and **Fig. 7N**). Collectively, these findings suggest that VTA-DA neurons hold a pivotal function in governing behavior in response to social and competing rewarding stimuli, and that *Shank3* mutation disrupts this function.

**Figure 7.**
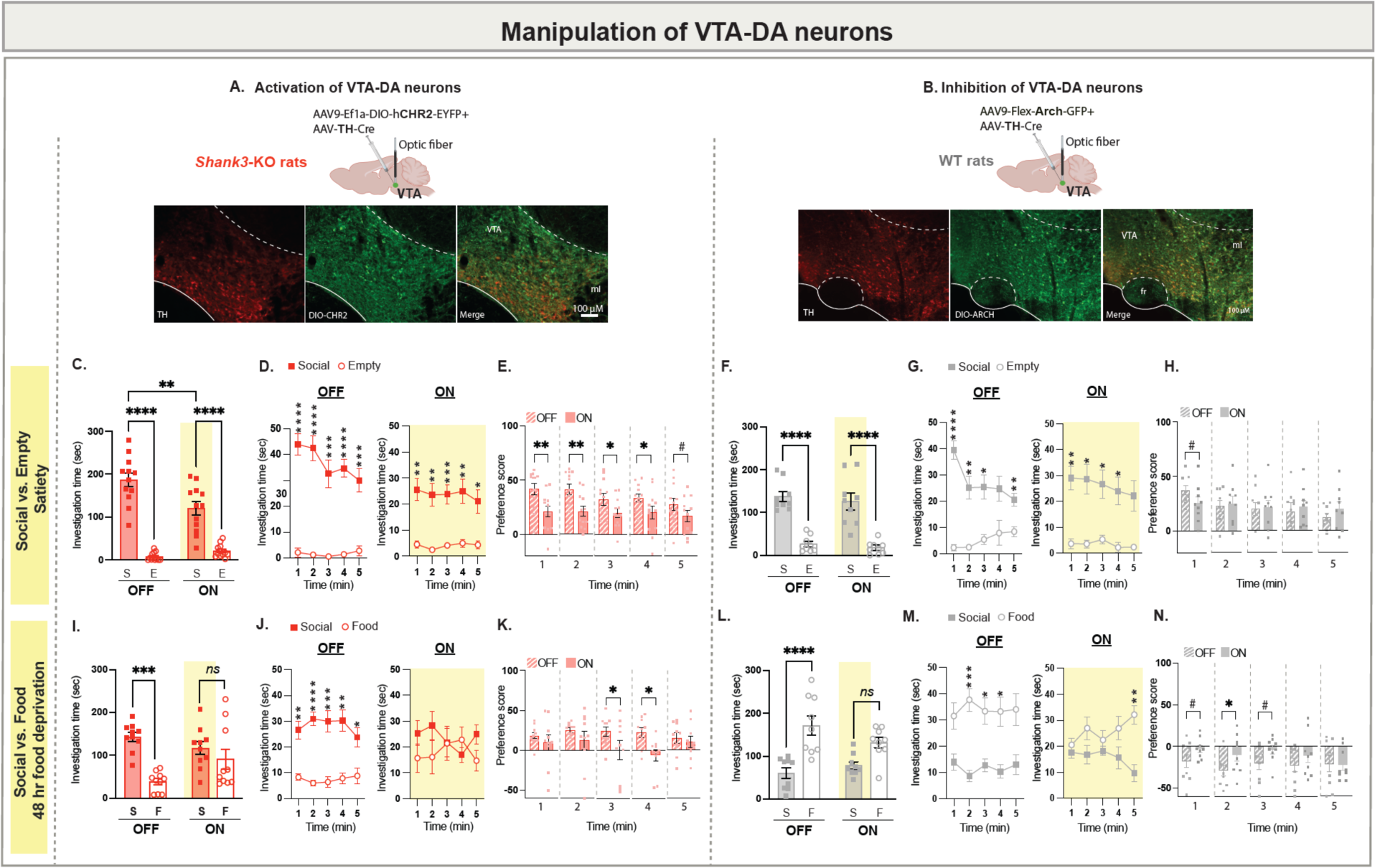
Optogenetic activation of VTA-DA neurons improves the atypical behavior observed in *Shank3*-deficient rats. **A.** A schematic of viral injection and optical fibers placement in the VTA of *Shank3*-KO rats (Top) and immunohistochemistry staining (bottom) of a sample VTA section showing TH expression (left panel, red), CHR2 (viral) expression (middle panel, green) and overlap (right panel). **B**. Same as A, but with ARCH expression in WT rats. 10X magnification; scale bar = 100 μm. **C.** Mean total investigation time of the social stimulus or the empty compartment during the non-activation (OFF) or activation (ON) of the VTA-DA neurons in *Shank3*-KO rats (n=12) when the rats approach the social stimulus. **D.** Mean total investigation time for the social stimulus and the empty compartment averaged in 1-minute intervals over the 5-minute test period in *Shank3*-KO rats. **E.** Differences in mean investigation time between the social stimulus and the empty compartment for each minute in *Shank3*-KO rats. **F-H.** Similar to C-E, but shows behavioral data during the non-inhibition (OFF) or inhibition (ON) of the VTA-DA neurons in WT rats (n=9) when they approach the social stimulus. **I-N**. Similar to C-H, but show behavioral data during the Social vs Food test at 48h of food deprivation in WT rats (KO, n=10; WT, n=9). *\*\*\*\*p*<.0001, *\*\*\*p*<.001, *\*\*p*<.01, *\*p*<.05, post hoc tests following the main effect. All error bars represent SEM. Detailed statistical data are provided as a Data file in Extended Data, Table 1.

### Optogenetic inhibition of VTA-GABA neurons normalizes the atypical behavior in *Shank3*-*KO* rats

Next, we sought to examine if optogenetic inhibition of VTA-GABA neurons could also normalize the atypical behavioral phenotype in *Shank3*-KO rats. To inhibit VTA-GABA neurons, we injected the rAAV-hVGAT1-Cre-WPRE-hGH polyA virus along with a Cre-dependent Arch; AAV9-FLEX-Arch-GFP virus, into the VTA (**Fig. 8A**). In parallel, we carried out a complementary experiment in WT rats to investigate whether activating VTA-GABA neurons could elicit an atypical behavioral phenotype similar to what is observed in *Shank3*-deficient rats. For this purpose, we injected the AAV-vGAT-Cre and AAV9-Ef1a-DIO-hChR2(E123T/T159C)-EYFP viruses into the VTA of WT rats (**Fig. 8B**). We found that inhibition of VTA-GABA neurons in *Shank3*-KO rats attenuated the atypical increase in interaction time with the social stimulus, which although was not captured by examining the overall investigation time (**Fig. 8C**), it was clearly detected when examining the behavioral pattern and the preference score across the 5 minutes of testing (**Fig. 8D** and **8E**). Activating VTA-GABA neurons in WT littermate rats during the Social vs. Empty task showed no significant effect on the rats’ total investigation time (**Fig. 8F**, light-ON), yet it influenced the pattern in which the rats investigated and expressed preference between the social and food compartments (**Fig. 8G** and **8H**), particularly during the first minute, where we noted a trend in the significance of the preference score (**Fig. 8H**). When tested on the Social vs. Food task, 48 hours after food deprivation where *Shank3*-KO rats still showed strong preference to the social stimulus despite food deprivation (**Fig. 8I** and **8J**, light OFF), we found that inhibition of VTA-GABA neurons in *Shank3*-KO rats during social interaction altered this preference (**Fig. 8I** and **8J**, light ON), mainly during the first and second minutes (**Fig. 8K**), causing their behavior to resemble that of their WT littermates (**Fig. 8L** and **8M**, light OFF), the latter which also changed following activation of VTA-GABA neurons (**Fig. 8M**, light ON), mainly during the first and second minutes of testing (**Fig. 8N**). These findings indicate that VTA-GABA neurons also play a crucial role in regulating behavior during social interaction, and that *Shank3* mutation interfere with this function too.

**Figure 8.**
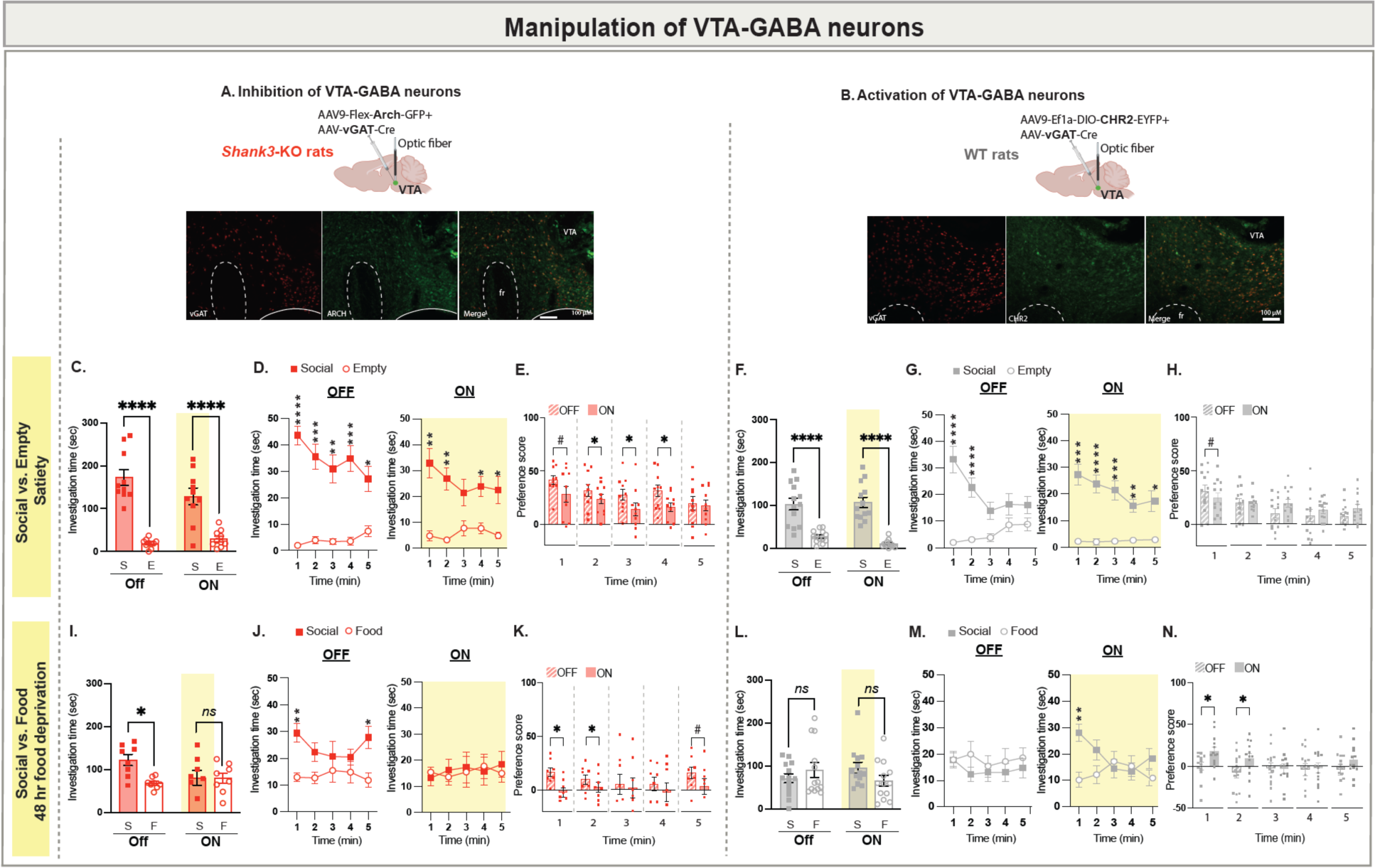
Optogenetic inhibition of VTA-GABA neurons improves the atypical behavior observed in *Shank3*-deficient rats. **A.** A schematic of viral injection and optical fibers placement in the VTA (Top) of *Shank3*-KO rats and immunohistochemistry staining (bottom) of a sample VTA section showing vGAT expression (left panel, red), CHR2 (viral) expression (middle panel, green) and overlap (right panel). **B**. Same as A, but with ARCH expression in WT rats. 10X magnification; scale bar = 100 μm. **C.** Mean total investigation time of the social stimulus or the empty compartment during the non-inhibition (OFF) or inhibition (ON) of the VTA-GABA neurons in *Shank3*-KO rats (n=10) when the rats approach the social stimulus. **D.** Mean total investigation time for the social stimulus and the empty compartment averaged in 1-minute intervals over the 5-minute test period in *Shank3*-KO rats. **E.** Differences in mean investigation time between the social stimulus and the empty compartment for each minute**. F-H.** Similar to C-E, but show behavioral data during the non-activation (OFF) or activation (ON) of the VTA-GABA neurons in WT rats (n=13) when they approach the social stimulus. **I-N**. Similar to C-H, but show behavioral data during the Social vs Food test at 48h of food deprivation (KO, n=8; WT, n=13). *\*\*\*\*p*<.0001, *\*\*\*p*<.001, *\*\*p*<.01, *\*p*<.05, post hoc tests following the main effect. All error bars represent SEM. Detailed statistical data are provided as a Data file in Extended Data, Table 1.

Our optogenetic experiments thus indicate that manipulating the activity of either DAergic or GABAergic neurons in the VTA enables the restoration of social behavior in *Shank3*-deficient rats, resembling the behavior observed in WT animals. Altogether, our findings suggest that *Shank3*-deficient rats exhibit impairment in processing social reward and deriving satisfaction from it, attributable to altered neuronal activity in the VTA

## Discussion

In this study, we employed in vivo recording techniques in *Shank3*-deficient rats to investigate how *Shank3* mutation affects the real-time function of the mesoaccumbens pathway during social and non-social interactions. Additionally, we used optogenetic tools to assess whether altering neural activity in the mesoaccumbens pathway could influence the rats’ behavior.

By conducting a thorough and impartial examination of behavioral traits, we identified that *Shank3*-deficient rats displayed an atypical increase in interaction with both newly introduced social and non-social stimuli when either was presented without any competing stimuli. This atypical behavior was particularly prominent in *Shank3*-KO rats and more pronounced during interaction with a social stimulus. Furthermore, when *Shank3*-deficient rats, deprived of food, were exposed to either a juvenile or a toy rat along with food, which held an elevated reward value during the state of food deprivation, an unusually heightened engagement with the social stimulus was observed in KO rats. This phenomenon was notable only in the presence of a juvenile rat stimulus, not a toy rat stimulus. Collectively, these findings illustrate the wide-ranging influence of *Shank3*-mutation on both the social and non-social interactions of the rats, as well as difficulty in their ability to adjust their behavior in response to alterations in the reward value of presented stimuli. Crucially, these effects seem to hinge on the specific context in which they unfold (i.e., presented with one vs. two rewarding social or non-social stimuli) and imply a deficiency in the rat’s ability to discern and respond to the positive value of social rewards, resulting in an atypical increase in interaction, that may not necessarily translate to a rewarding experience.

In this context, it is important to emphasize the persistent misconception in the preclinical autism research community that impaired social interactions in ASD invariably lead to a reduction in overall social engagement. Social deficits in ASD and associated neurodevelopmental disorders are highly heterogeneous. Indeed, PMS serves as an exemplar of the unique clinical manifestation of ASD, especially when juxtaposed with idiopathic ASD, which is inherently characterized by its high heterogeneity. This divergence is observable in several key aspects, including the restricted and repetitive behaviors^71^, sensory symptoms^72^, functional MRI changes in response to social versus non-social sounds^73^, and social deficits, which in contrast to the presentation in idiopathic ASD, do not always manifest as social aversion^74,75^. Another prevailing but mistaken assumption in the field is that behavioral deficits in animal models should perfectly replicate those observed in humans with equivalent genetic alterations. This topic was recently discussed by Silverman et al., who also underscored the importance of the model’s construct validity and the utilization of reproducible and validated outcome measures^76^. This approach, as opposed to trying to artificially force the model into a human-like phenotype, optimally advances our goal of deepening our understanding of the mechanistic effects of mutations associated with neurodevelopmental disorders.

The increased social interaction we observed in *Shank3*-deficient rats and their inability to adjust behavior according to the rewarding value of the presented stimulus, prompted us to test the hypothesis that deficits in the mesoaccumbens reward pathway could be a contributing factor. The role of VTA-DA neurons in learning and motivation, as well as in reward processing has been well-established in both clinical and preclinical studies^10,11,16–19^. Specifically, studies in mice have highlighted the critical involvement of VTA-DA neurons in the processing of social rewards, with increased activity noted during social interactions^46,47,66,68^. Consistent with these observations, our findings demonstrated, for the first time, increased activity in VTA-DA neurons also in rats (WT rats) when they engaged with a social stimulus. However, this pattern of activity was not observed in *Shank3*-deficient rats, despite their increased interaction with the social stimulus. Additionally, when compared to their WT littermates, *Shank3*-deficient rats exhibited increased GABAergic activity during social interaction. These deficiencies are in alignment with earlier findings from *in vitro* and *in vivo* single unit recordings conducted in mice, which demonstrated that reducing Shank3 levels in the VTA during early postnatal development led to decreased activity in DA neurons and increased activity in GABA neurons, which was linked to an impaired maturation of excitatory synapses on these neurons^47^. The question of whether the lack of maturation in excitatory synapses on VTA neurons accounts for the impaired VTA neural activity during social interaction in the rat model remains a subject for future investigations. In this context, it is important to emphasize that the activity of DAergic and GABAergic neurons of the VTA is modulated not only by excitatory but also inhibitory inputs from various brain regions, including inputs from the BNST, LH, NAc, MPOA^17,77–79^. Activation of these inputs was demonstrated to induce a pro-reward behavior, increased reward consumption, and/or reduced anxiety^80^. In the realm of social reward, activation of the LH–VTA GABA pathway has been demonstrated to enhances social interaction. This occurs via suppression of VTA-GABA neurons and facilitation of DA release in the nucleus accumbens^81^. A recent study in rats has shown that inhibition of CeA projections onto VTA-GABA neurons disrupts the maintenance, but not the initiation, of social interaction, potentially due to disinhibition of DA neurons in the VTA^82^. These findings together propose a model in which local VTA-GABA circuits shape social behavior. They also imply that the influence of a deficiency in Shank3 on VTA-DA neurons may occur indirectly through its impact on the intricate nature of the VTA-GABA population. Therefore, future investigations should further explore the impact of *Shank3*-deficiency on VTA local circuits to advance our comprehension of the pathophysiology related to *Shank3*-deficiencies. Additionally, considering the diverse nature of VTA neural populations and the intricate network of inputs and outputs within the VTA^17,77–79^, as well as their distinct roles in various behaviors, including feeding, learning and motivation, and social behavior^80^, it is imperative for future studies to dissect the specific influence of *Shank3* in particular VTA pathways, encompassing both social and non-social behaviors. An initial step towards addressing this broader question could involve mapping the expression of *Shank3* in subpopulations of neurons and specific VTA inputs and outputs.

Extensive research has established the pivotal role of VTA-DA to NAc projecting neurons and DA release in the NAc in motivated behaviors, reward processing, and social behvaior^10,11,16–19^. Moreover, VTA-GABA neurons were shown to inhibit the activation of VTA-DA neurons potentially perturbing DA release in the NAc^83^. Pertaining to social behavior, studies in rats have demonstrated that variations in dopamine levels in both the dorsal and ventral striatum correspond with episodes of social interaction^84,85^. Moreover, a prior study in mice has elucidated that social interaction leads to heightened activity, particularly in VTA-DA neurons projecting to the NAc^66^.

These observations align with our findings in WT rats, illustrating an increase in DA release in the NAc during social interaction. This increase likely plays a crucial role in encoding the rewarding value of social interactions, leading to a gradual decrease in motivation for further social engagement once that reward is registered and satisfaction is achieved. Our findings in *Shank3*-KO rats, which highlight that reduced DA release in the NAc and compromised activity in both VTA-DA and VTA-GABA neurons are linked to an atypical increase in social interaction, suggest that these rats experience a deficiency in processing the rewarding value of social interaction. As a consequence of this deficiency, there is reduced satisfaction from social interactions, prompting them to pursue more of these interactions without properly adapting their behavior to other rewarding stimuli in their surroundings.

Overall, our findings demonstrate that Shank3 plays a critical role in modulating the activity of VTA neurons and the VAT-NAc pathway, which plays pivotal role during reward processing of social stimuli. In support of this argument, our optogenetic interventions provided evidence of a causal relationship between the observed deficits in neural activity and the atypical social interactions in the *Shank3* rat model. Additionally, our discovery that this atypical interaction could be ameliorated by augmenting the activity of VTA-DA neurons or mitigating the excessive activity of VTA-GABA neurons in *Shank3*-deficient rats during adulthood implies that there may still be opportunities to intervene in the VTA pathway and potentially improve social behavioral deficits associated with *Shank3-*deficiencies.

## Supporting information

Extended_data_Table_1_Statistics

## Acknowledgments

This work was supported by National Institute of Mental Health (Grant No. R01MH116108 [to H.H.N]) and a Seaver Foundation fellowship (to M.B and H.H.N). S.W was supported by the ISF-NSFC joint research program (Grant No. 3459/20), the Israel Science Foundation (Grant Nos. 1361/17 and 2220/22), the Ministry of Science, Technology and Space of Israel (Grant No. 3-12068), the Ministry of Health of Israel (Grant No. 3-18380), the German Research Foundation (DFG) (Grant Nos. GR 3619/16-1 and SH 752/2-1), the Congressionally Directed Medical Research Programs (CDMRP) (Grant No. AR210005) and the United States-Israel Binational Science Foundation (BSF) (Grant No. 2019186, S.W and H.H.N). We thank Dr. Arthur Godino (Dr. Eric Nestler’s Lab at Mount Sinai) for sharing fiber photometry equipment and Dr. Matthew Perkins (Dr. Ivan De Araujo’s lab at Mount Sinai) for assisting with optogenetic set-up and resources. The authors would like to thank the Microscopy and Advanced Bioimaging CoRE core at the Icahn School of Medicine for their guidance on imaging. Parts of Figures 1-8 and Extended Data Figures 9 and 10 were created with https://BioRender.com.

## Author contributions

M.B and H.H.N conceptualized and designed the study. M.B performed all experiments and acquired all imaging data. K.T.R. helped on the behavioral analysis and contributed to the conceptualization of the study and the manuscript preparation. S.N and S.W provided support on the open-source behavioral analysis software and manuscript preparation. M.B and H.H.N, interpreted the data, prepared figures, and wrote the manuscript. All authors read and approved the final manuscript.

---

**Extended Data Figure 1.**
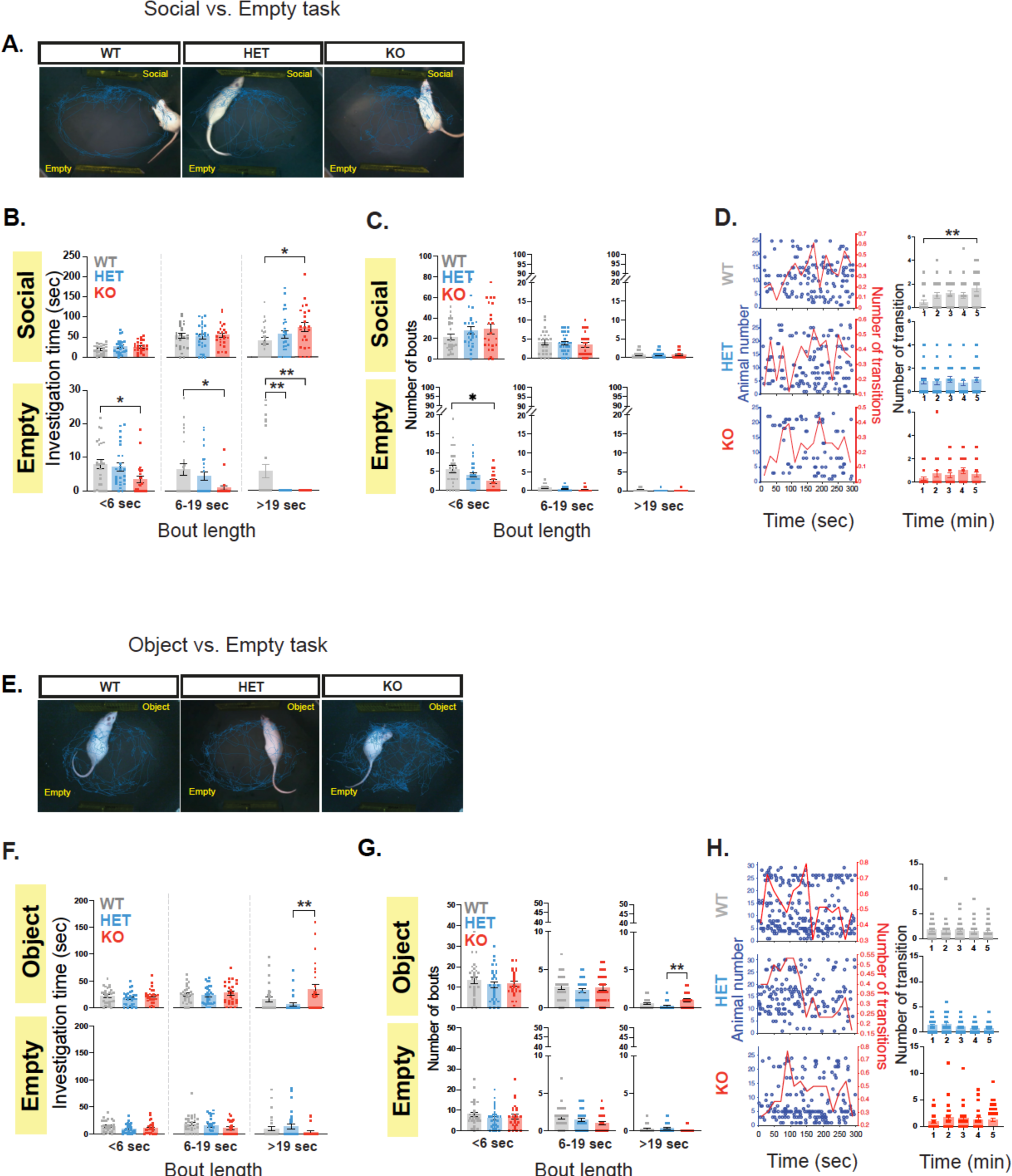
**A.** A representative trace (from one rat per genotype) during the Social vs. Empty task. **B.** Mean combined duration of short investigation bouts (<6 sec), medium length bouts (6-19 sec), and long investigation bouts (>19 sec) during the Social vs. Empty task during investigation of the social-containing (top) or empty compartment (bottom). **C.** Mean combined number of investigation bouts (<6 sec), medium length bouts (6-19 sec), and long investigation bouts (>19 sec) during the Social vs. Empty task during investigation of the social-containing (top) or empty compartment (bottom). **D**. Left, Transitions between the two compartments across time during the Social vs. Empty task. Each punctum denotes the beginning of investigation of a new stimulus, and each row represents a single subject. The mean rate (using 20-s bins) is denoted by the red line (right red y-axis). Right, Mean pooled number of transitions between the two compartments (social or empty) over the 5-minute test period. WT, n=25; HET, n=26; KO, n=23. **E-H.** Similar to A-D, but shows behavior during the Object vs. Empty task. WT, n=29; HET, n=30; KO, n=26. *\*\*p*<.01, *\*p*<.05, post hoc tests following the main effect. All error bars represent SEM. Detailed statistical data are provided as a Data file in Extended Data, Table 1.

**Extended Data Figure 2.**
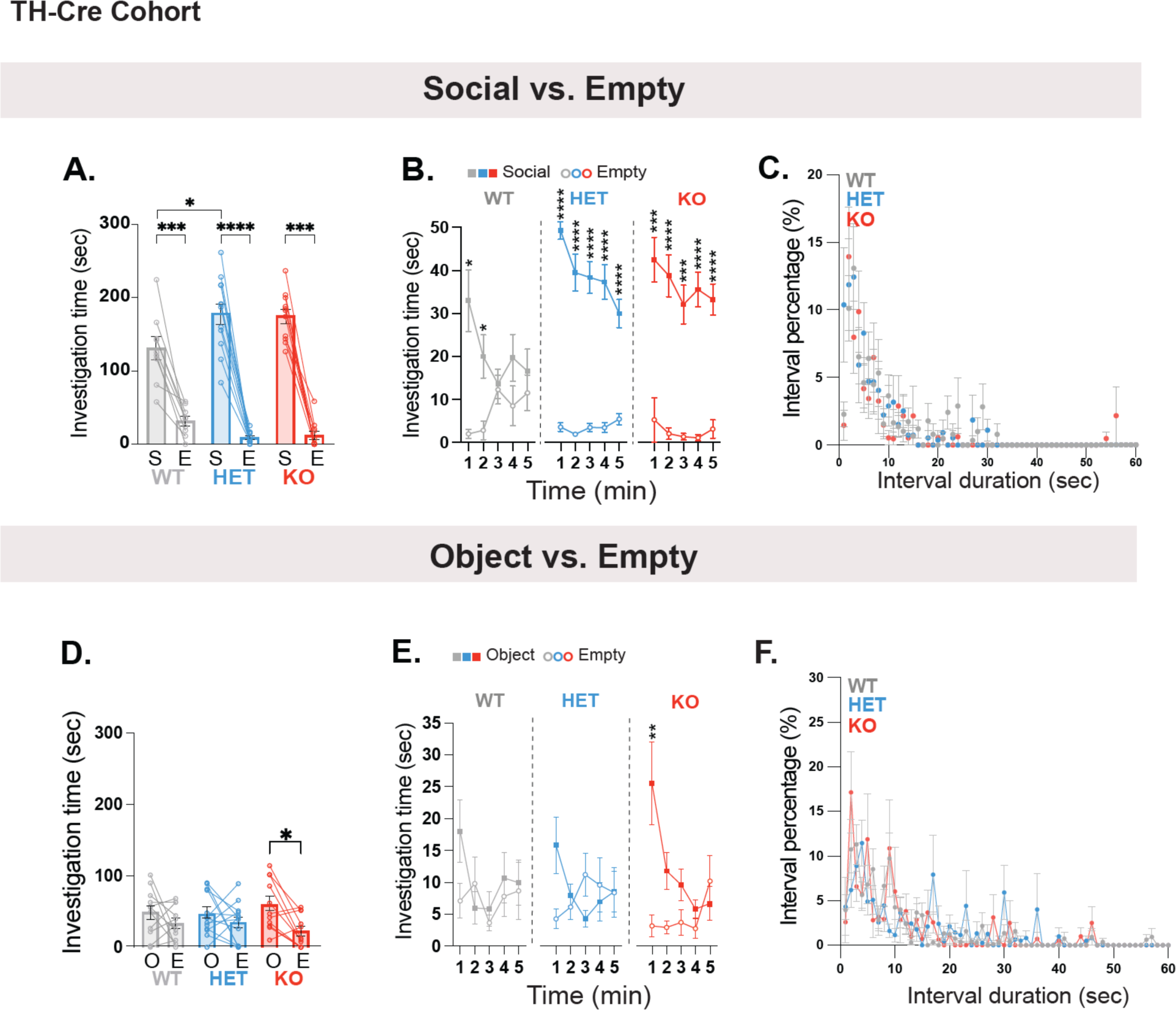
**A.** Mean total investigation time of the social stimulus or the empty compartment for the cohort of rats injected with TH-Cre and DIO-GCamp6 viruses, during testing on the Social vs. Empty task. **B.** Mean total investigation time for the social stimulus and the empty compartment averaged in 1-minute intervals over the 5-minute test period. WT, n=9; HET, n=12; KO, n=11. **C.** Mean percentage of each interval length (using 10 sec bins) during the 5-minutes of testing on the Social vs. Empty task. Intervals duration represent the time it takes an animal to transition from one compartment to another. **D-F.** Similar to A-C, but shows behavior during the Object vs. Empty task. WT, n=11; HET, n=13; KO, n=11. *\*\*\*\*p*<.0001, *\*\*\*p*<.001, *\*\*p*<.01, *\*p*<.05, post hoc tests following the main effect. All error bars represent SEM. Detailed statistical data are provided as a Data file in Extended Data, Table 1.

**Extended Data Figure 3.**
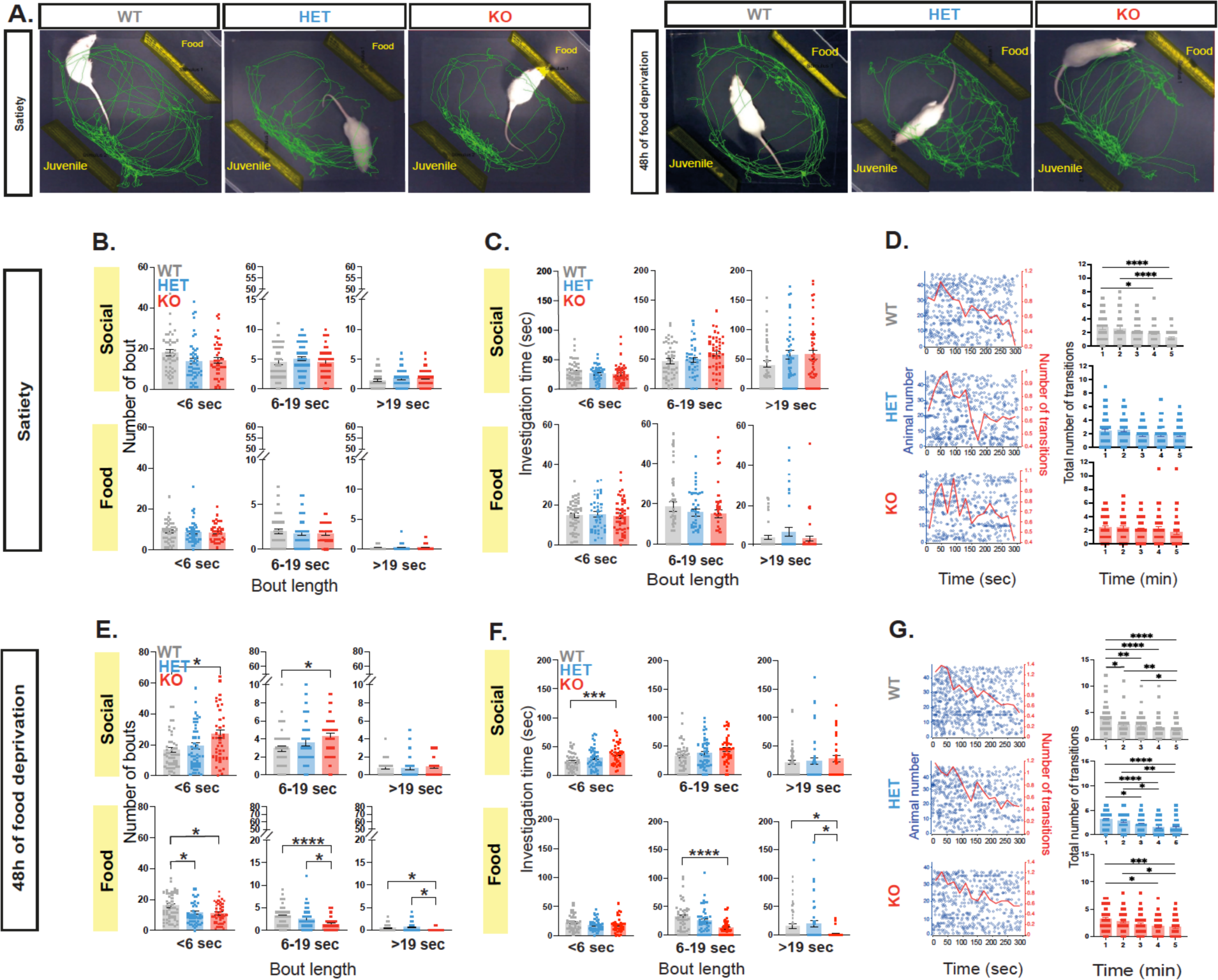
**A.** A representative trace (from one rat per genotype) during the Social vs. Food task, at satiety (left) and after 48 hours of food deprivation (right). **B.** Mean combined number of investigation bouts (<6 sec), medium length bouts (6-19 sec), and long investigation bouts (>19 sec) during the Social vs. Food task at satiety during investigation of the social-containing (top) or food compartment (bottom).**C.** Mean combined duration of short investigation bouts (<6 sec), medium length bouts (6-19 sec), and long investigation bouts (>19 sec) during the Social vs. Food task during investigation of the social-containing (top) or food compartment (bottom). **D**. Left, Transitions between the two compartments across time during the Social vs. Food task. Each punctum denotes the beginning of investigation of a new stimulus, and each row represents a single subject. The mean rate (using 20-s bins) is denoted by the red line (right red y-axis). Right, Mean pooled number of transitions between the two compartments (social or food) over the 5-minute test period. **E-G**. Similar to B-D, but shows behavior during the Social vs Food task after 48h of food deprivation. WT, n=47; HET, n=47; KO, n=41. *\*\*\*\*p*<.0001, *\*p*<.05, post hoc tests following the main effect. All error bars represent SEM. Detailed statistical data are provided as a Data file in Extended Data, Table 1.

**Extended Data Figure 4.**
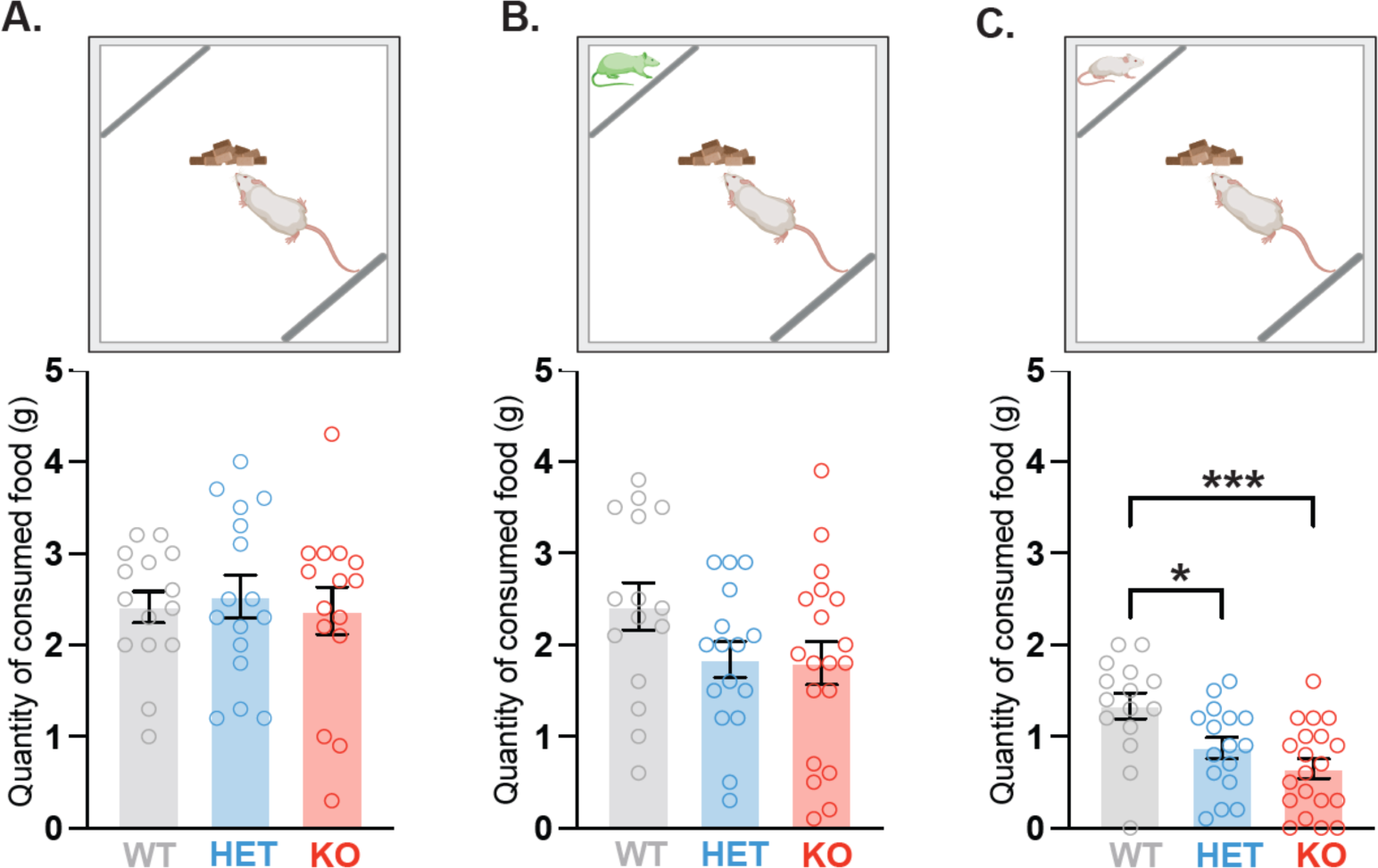
**A.** Outline for the Free Food Intake Test (Top). Graph depicting the average food intake during the 5-minute test period where no stimulus is presented (Bottom). **B.** Similar to A, but shows food intake during the Object vs. Empty task. **C.** Similar to A, but shows food intake during the Social vs. Empty task. WT, n=15; HET, n=16; KO, n=15. All error bars represent SEM. *\*\*\*p*<.001, *\*p*<.05, post hoc tests following the main effect. Detailed statistical data are provided as a Data file in Extended Data, Table 1.

**Extended Data Figure 5.**
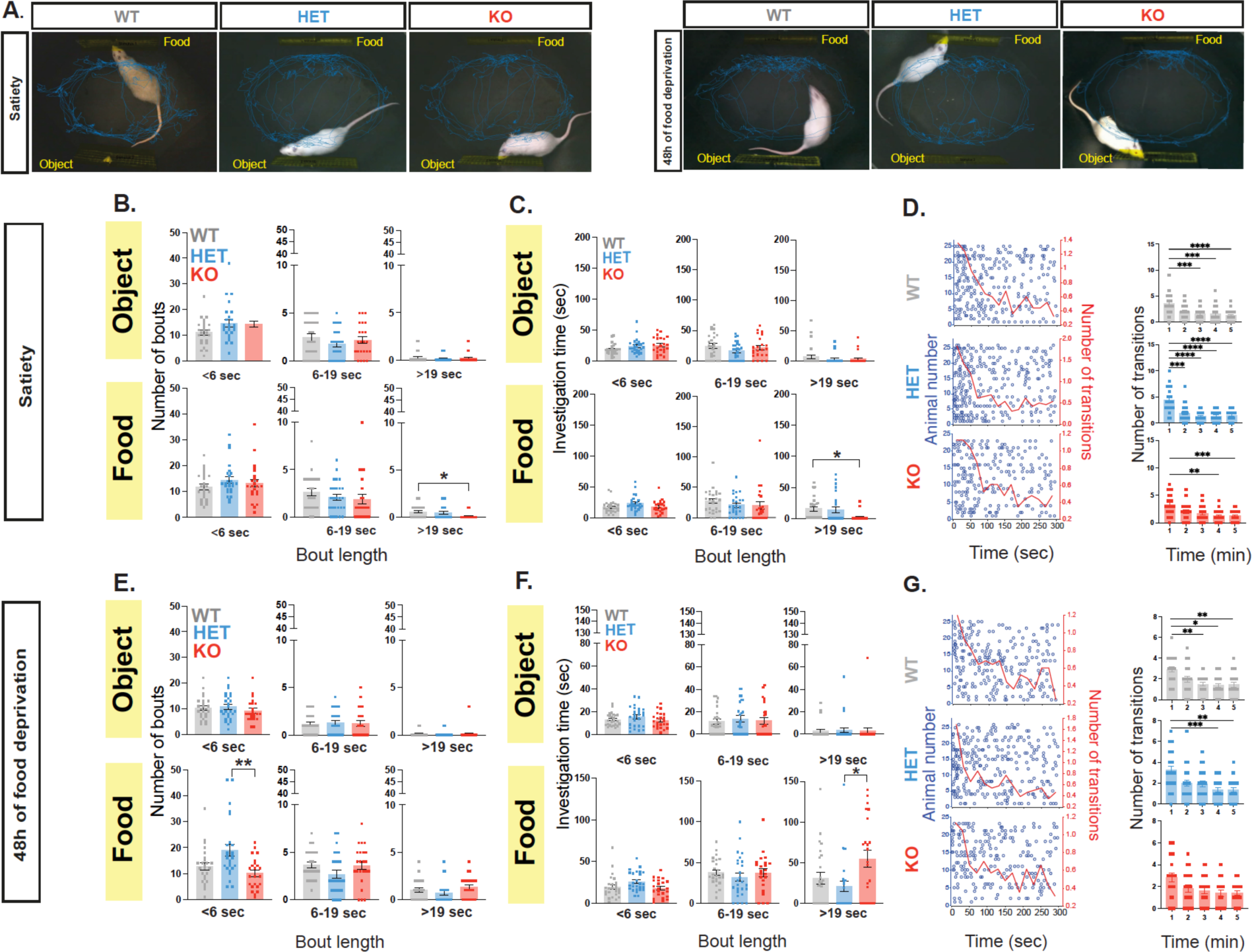
**A.** A representative trace (from one rat per genotype) during the Object vs. Food task, at satiety (left) and after 48 hours of food deprivation (right). **B.** Mean combined number of investigation bouts (<6 sec), medium length bouts (6-19 sec), and long investigation bouts (>19 sec) during the Object vs. Food task during investigation of the moving object-containing (top) or food compartment (bottom) at satiety. **C.** Mean combined duration of short investigation bouts (<6 sec), medium length bouts (6-19 sec), and long investigation bouts (>19 sec) during the Object vs. Food task during investigation of the object-containing (top) or food compartment (bottom). **D**. Left, Transitions between the two compartments across time during the Object vs. Food task. Each punctum denotes the beginning of investigation of a new stimulus, and each row represents a single subject. The mean rate (using 20-s bins) is denoted by the red line (right red y-axis). Right, Mean pooled number of transitions between the two compartments (object or food) over the 5-minute test period. **E-G**. Similar to B-D, but shows behavior during the Object vs Food task after 48 hours of food deprivation. WT, n=25; HET, n=26; KO, n=23. *\*p*<.05, post hoc tests following the main effect. All error bars represent SEM. Detailed statistical data are provided as a Data file in Extended Data, Table 1.

**Extended Data Figure 6.**
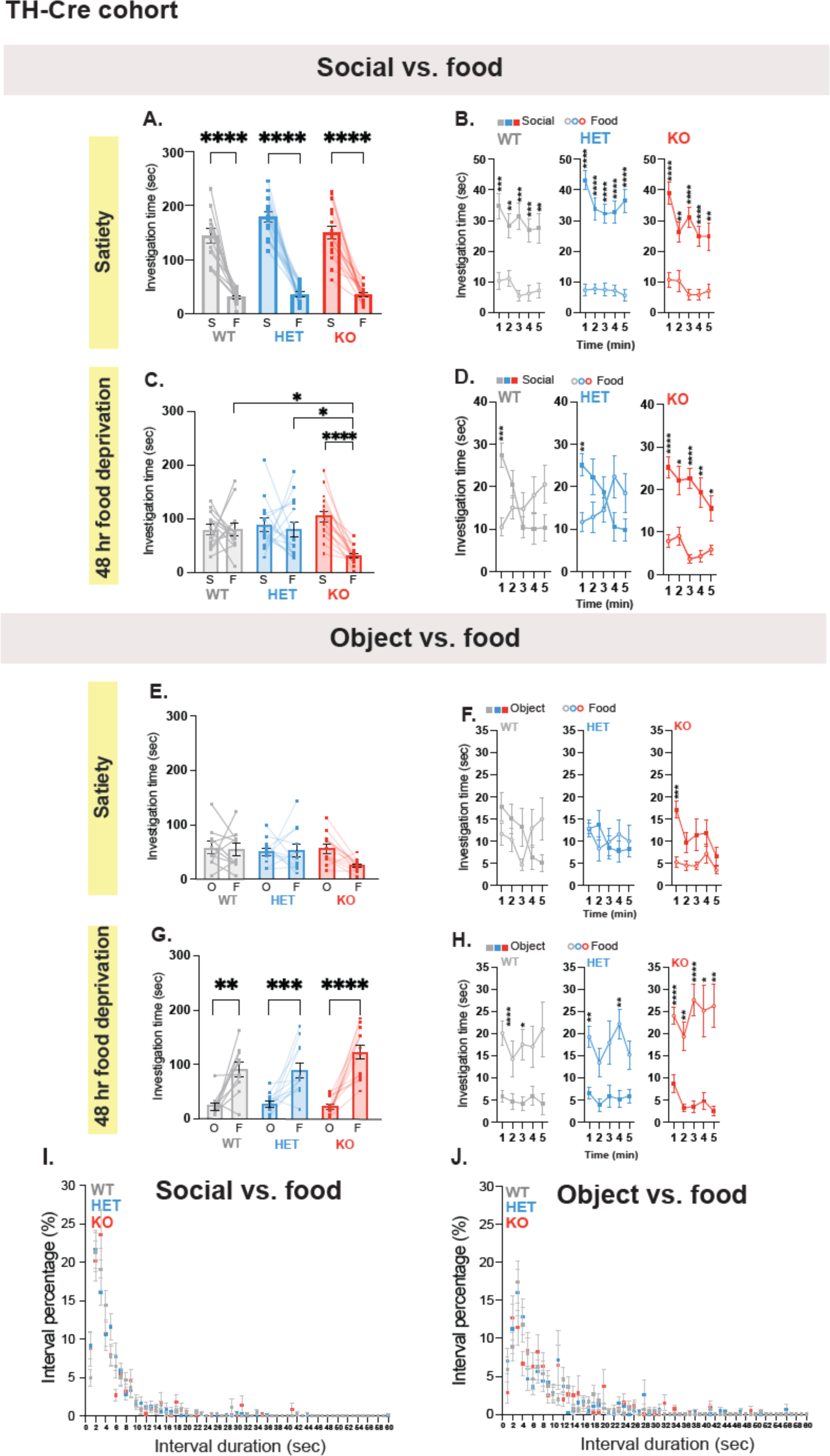
**A.** Mean total investigation time of the social stimulus or the food compartment at satiety for the cohort of rats injected with TH-Cre and DIO-GCamp6 viruses, during testing on the Social vs. Food task. **B.** Mean total investigation time for the social stimulus and the food compartment at satiety, averaged in 1-minute intervals over the 5-minute test period. **C and D.** Similar to A and B, but shows behavior after 48h of food deprivation. WT, n=13; HET, n=15; KO, n=16. **E-H.** Similar to A-D, but shows behavior during the Object vs Food task. WT, n=10; HET, n=12; KO, n=13. *\*\*\*\*p*<.0001, *\*\*\*p*<.001, *\*\*p*<.01, *\*p*<.05, post hoc tests following the main effect. All error bars represent SEM. Detailed statistical data are provided as a Data file. **I** and **J**. Mean percentage of each interval length (using 10 sec bins) during the 5-minutes of testing on the Social vs. Food task (I) or Object vs Food task (J). Intervals duration represent the time it takes an animal to transition from one compartment to another. Detailed statistical data are provided as a Data file in Extended Data, Table 1.

**Extended Data Figure 7.**
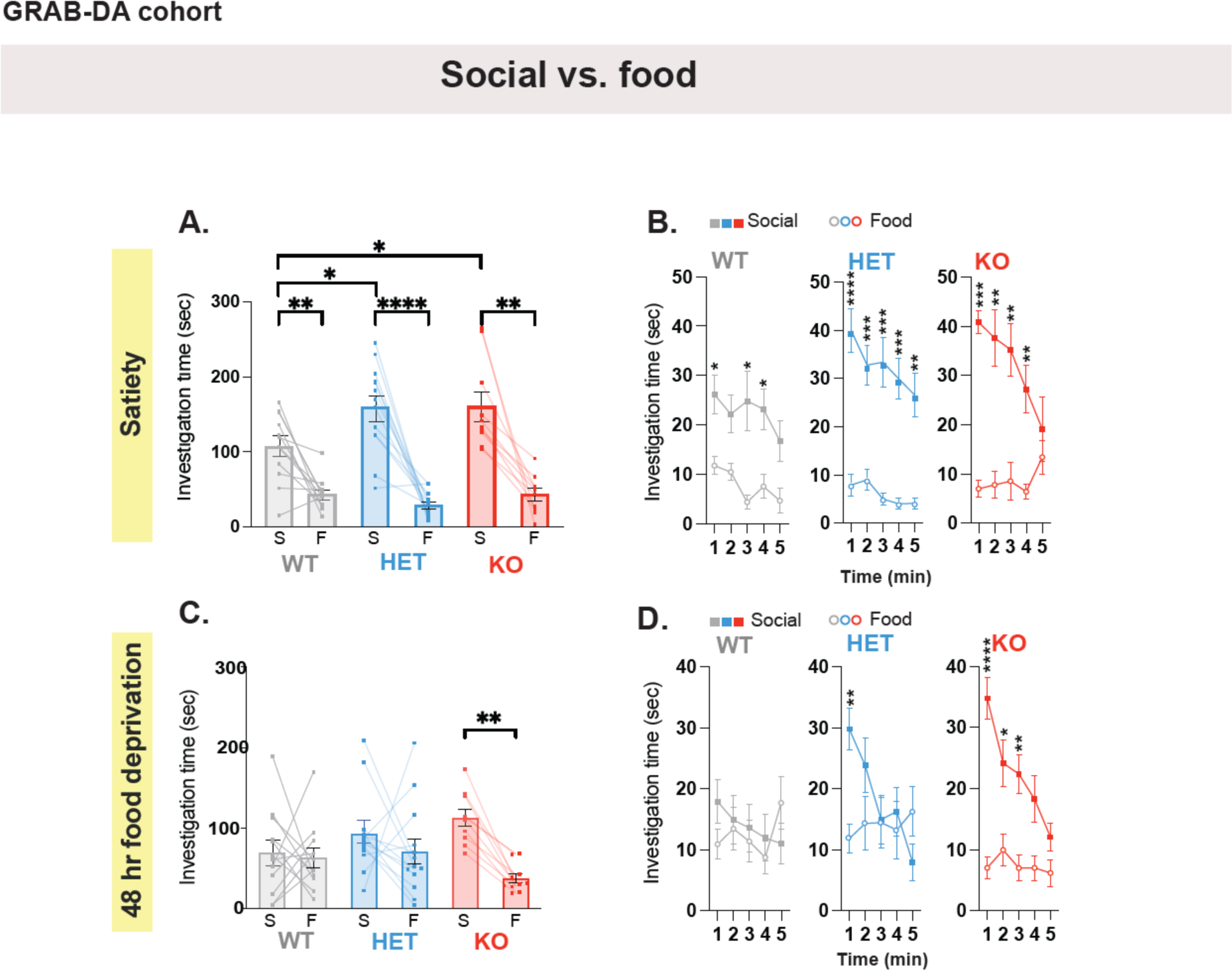
**A.** Mean total investigation time of the social stimulus or the food compartment at satiety for the cohort of rats injected with GRAB^DA,^ during testing on the Social vs. Food task. **B.** Mean total investigation time for the social stimulus and the food compartment averaged in 1-minute intervals over the 5-minute test period. **C** and **D.** Similar to A and B respectively, but show behavior after 48 hours of food deprivation. WT, n=12; HET, n=13; KO, n=10. *\*\*\*\*p*<.0001, *\*\*\*p*<.001, *\*\*p*<.01, *\*p*<.05, post hoc tests following the main effect. All error bars represent SEM. Detailed statistical data are provided as a Data file in Extended Data, Table 1.

**Extended Data Figure 8.**
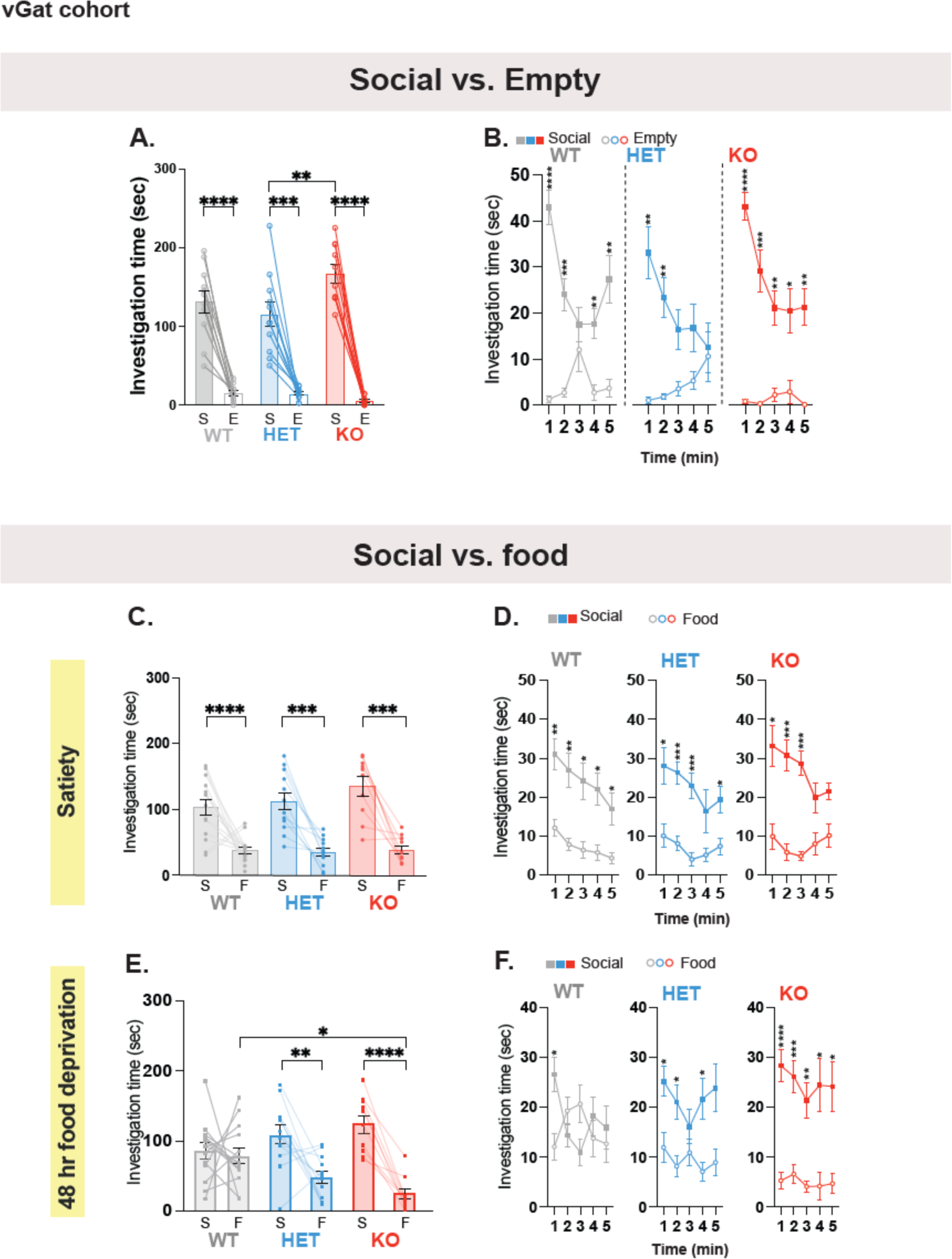
**A.** Mean total investigation time of the social stimulus or the empty compartment for the cohort of rats injected with vGAT-Cre and DIO-GCamp6 viruses, during testing on the Social vs. Empty task. **B.** Mean total investigation time for the social stimulus and the empty compartment averaged in 1-minute intervals over the 5-minute test period. WT, n=11; HET, n=11; KO, n=10. **C** and **D.** Similar to A and B, respectively, but show behavior during the Object vs Food task at satiety (C) and after 48 hours of food deprivation (D). WT, n=14; HET, n=12; KO, n=10. *\*\*\*\*p*<.0001, *\*\*\*p*<.001, *\*\*p*<.01, *\*p*<.05, post hoc tests following the main effect. All error bars represent SEM. Detailed statistical data are provided as a Data file in Extended Data, Table 1.

**Extended Data Figure 9.**
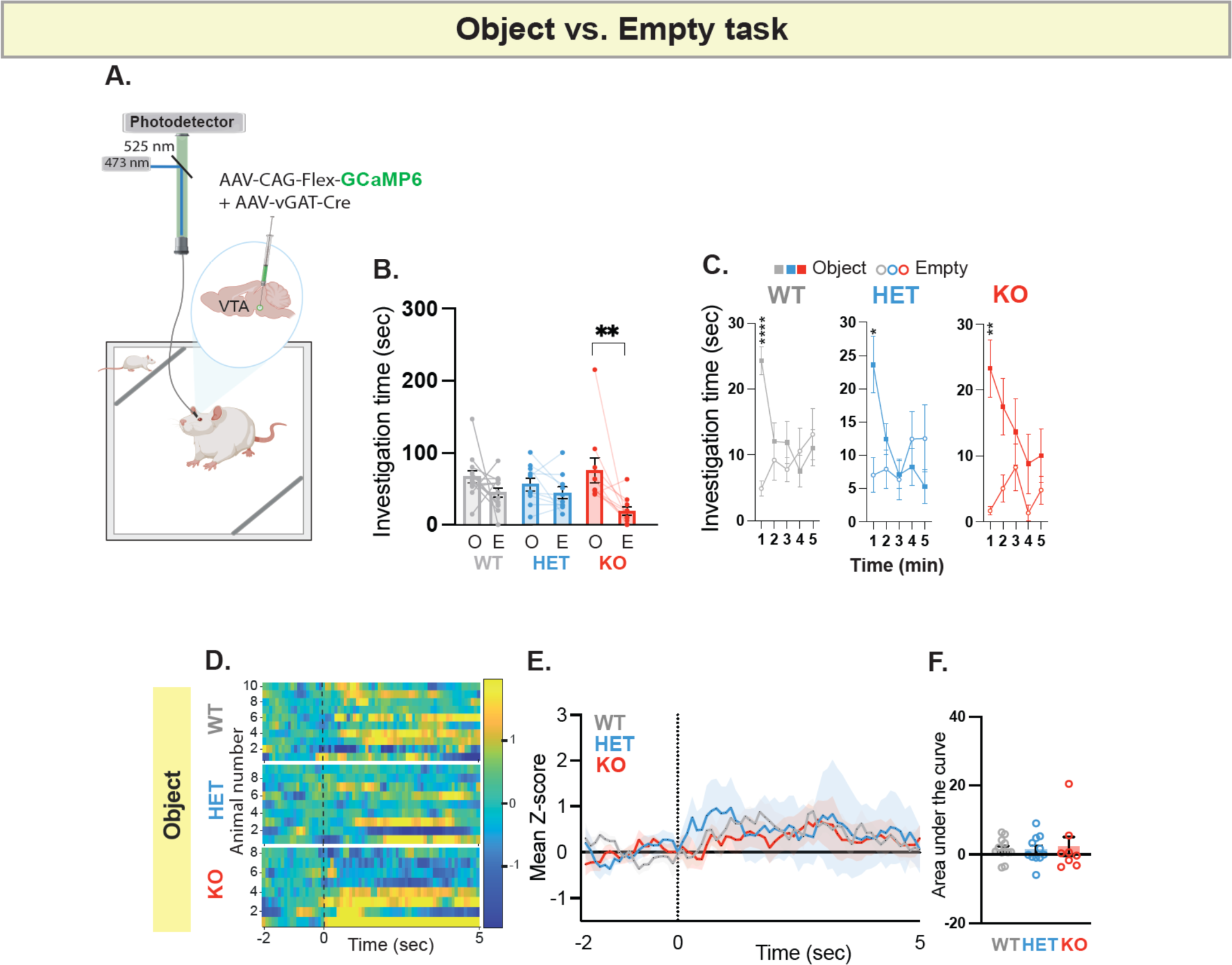
**A.** A schematic of viral injection and fiber photometry setup in the VTA to record neural activity of VTA-GABAergic neurons. **B.** Mean total investigation time of the object stimulus or the empty compartment for the cohort of rats injected with vGAT-Cre and DIO-GCamp6 viruses, during testing on the Object vs. Empty task. **C.** Mean total investigation time for the object stimulus and the empty compartment averaged in 1-minute intervals over the 5-minute test period. **D.** Heat maps illustrate the change in fluorescence signals (dF/F) from 2 seconds before to 5 seconds after each interaction bout with the object. Each row corresponds to one animal and comprises an average of signals from all interaction bouts during the 5-minute testing period. **E.** Average standardized VTA-vGAT photometry responses aligned to investigation onset of the object stimulus. **F.** The area under the curve, calculated from average standardized traces in (E). WT, n=13; HET, n=10; KO, n=10. *\*\*\*\*p*<.0001, *\*\*p*<.01, *\*p*<.05, post hoc tests following the main effect. All error bars represent SEM. Detailed statistical data are provided as a Data file in Extended Data, Table 1.

**Extended Data Figure 10.**
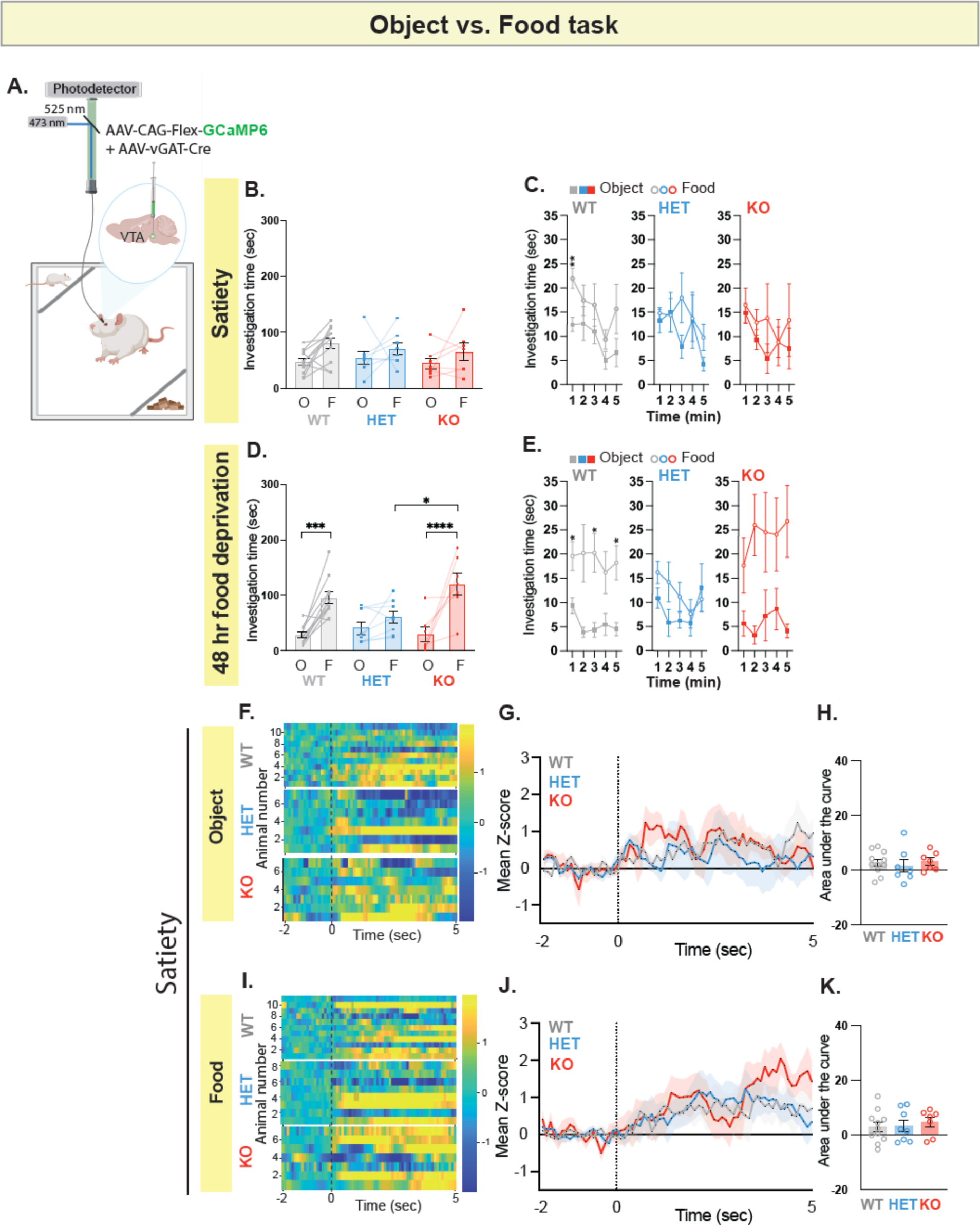
**A.** A schematic of viral injection and fiber photometry setup in the VTA to record neural activity of VTA-GABAergic neurons. **B.** Mean total investigation time of the object stimulus or the food compartment at satiety for the cohort of rats injected with vGAT-Cre and DIO-GCamp6 viruses, during testing on the Social vs. Food task. **C.** Mean total investigation time for the object stimulus and the food compartment averaged in 1-minute intervals over the 5-minute test period. **D** and **E.** Similar to B and C, respectively, but show behavior after 48 hours of food deprivation. **F.** Heat maps illustrate the change in fluorescence signals (dF/F) from 2 seconds before to 5 seconds after each interaction bout with the object. Each row corresponds to one animal and comprises an average of signals from all interaction bouts during the 5-minute testing period. **G.** Average standardized VTA-vGAT photometry responses aligned to investigation onset of the object stimulus. **H.** The area under the curve, calculated from average standardized traces in (G). **I-K.** Similar to F-H, but show fiber photometry signal for the food investigation. WT, n=11; HET, n=8; KO, n=7. *\*\*\*\*p*<.0001, *\*\*p*<.01, *\*p*<.05, post hoc tests following the main effect. All error bars represent SEM. Detailed statistical data are provided as a Data file in Extended Data, Table 1.

## References

1. American Psychiatric Association, D. (2013). American Psychiatric AssociationDiagnostic and Statistical Manual of Mental Disorders.

2. Assaf, M., Hyatt, C.J., Wong, C.G., Johnson, M.R., Schultz, R.T., Hendler, T., and Pearlson, G.D. (2013). Mentalizing and motivation neural function during social interactions in autism spectrum disorders. Neuroimage Clin 3, 321–331. 10.1016/j.nicl.2013.09.005.

3. Baumeister, S., Moessnang, C., Bast, N., Hohmann, S., Aggensteiner, P., Kaiser, A., Tillmann, J., Goyard, D., Charman, T., Ambrosino, S., et al. (2023). Processing of social and monetary rewards in autism spectrum disorders. Br J Psychiatry 222, 100–111. 10.1192/bjp.2022.157.

4. Chevallier, C., Kohls, G., Troiani, V., Brodkin, E.S., and Schultz, R.T. (2012). The social motivation theory of autism. Trends in cognitive sciences 16, 231–239. 10.1016/j.tics.2012.02.007.

5. Delmonte, S., Balsters, J.H., McGrath, J., Fitzgerald, J., Brennan, S., Fagan, A.J., and Gallagher, L. (2012). Social and monetary reward processing in autism spectrum disorders. Molecular autism 3, 7. 10.1186/2040-2392-3-7.

6. Dichter, G.S., Felder, J.N., Green, S.R., Rittenberg, A.M., Sasson, N.J., and Bodfish, J.W. (2012). Reward circuitry function in autism spectrum disorders. Soc Cogn Affect Neurosci 7, 160–172. 10.1093/scan/nsq095.

7. Kohls, G., Schulte-Ruther, M., Nehrkorn, B., Muller, K., Fink, G.R., Kamp-Becker, I., Herpertz-Dahlmann, B., Schultz, R.T., and Konrad, K. (2013). Reward system dysfunction in autism spectrum disorders. Soc Cogn Affect Neurosci 8, 565–572. 10.1093/scan/nss033.

8. Scott-Van Zeeland, A.A., Dapretto, M., Ghahremani, D.G., Poldrack, R.A., and Bookheimer, S.Y. (2010). Reward processing in autism. Autism Res 3, 53–67. 10.1002/aur.122.

9. Supekar, K., Kochalka, J., Schaer, M., Wakeman, H., Qin, S., Padmanabhan, A., and Menon, V. (2018). Deficits in mesolimbic reward pathway underlie social interaction impairments in children with autism. Brain 141, 2795–2805. 10.1093/brain/awy191.

10. Berridge, K.C., and Kringelbach, M.L. (2015). Pleasure systems in the brain. Neuron 86, 646–664. 10.1016/j.neuron.2015.02.018.

11. Berridge, K.C., and Robinson, T.E. (1998). What is the role of dopamine in reward: hedonic impact, reward learning, or incentive salience? Brain Res Brain Res Rev 28, 309–369. 10.1016/s0165-0173(98)00019-8.

12. Kalivas, P.W., and Nakamura, M. (1999). Neural systems for behavioral activation and reward. Curr Opin Neurobiol 9, 223–227. 10.1016/s0959-4388(99)80031-2.

13. Lammel, S., Lim, B.K., and Malenka, R.C. (2014). Reward and aversion in a heterogeneous midbrain dopamine system. Neuropharmacology 76 Pt B, 351–359. 10.1016/j.neuropharm.2013.03.019.

14. Lammel, S., Lim, B.K., Ran, C., Huang, K.W., Betley, M.J., Tye, K.M., Deisseroth, K., and Malenka, R.C. (2012). Input-specific control of reward and aversion in the ventral tegmental area. Nature 491, 212–217. 10.1038/nature11527.

15. Mirenowicz, J., and Schultz, W. (1996). Preferential activation of midbrain dopamine neurons by appetitive rather than aversive stimuli. Nature 379, 449–451. 10.1038/379449a0.

16. Schultz, W. (2016). Dopamine reward prediction-error signalling: a two-component response. Nature reviews. Neuroscience 17, 183–195. 10.1038/nrn.2015.26.

17. Watabe-Uchida, M., Eshel, N., and Uchida, N. (2017). Neural Circuitry of Reward Prediction Error. Annu Rev Neurosci 40, 373–394. 10.1146/annurev-neuro-072116-031109.

18. Wise, R.A. (2004). Dopamine, learning and motivation. Nature reviews. Neuroscience 5, 483–494. 10.1038/nrn1406.

19. Wise, R.A., and Rompre, P.P. (1989). Brain dopamine and reward. Annu Rev Psychol 40, 191–225. 10.1146/annurev.ps.40.020189.001203.

20. Brown, M.T., Tan, K.R., O’Connor, E.C., Nikonenko, I., Muller, D., and Luscher, C. (2012). Ventral tegmental area GABA projections pause accumbal cholinergic interneurons to enhance associative learning. Nature 492, 452–456. 10.1038/nature11657.

21. Morales, M., and Margolis, E.B. (2017). Ventral tegmental area: cellular heterogeneity, connectivity and behaviour. Nature reviews. Neuroscience 18, 73–85. 10.1038/nrn.2016.165.

22. Qi, J., Zhang, S., Wang, H.L., Barker, D.J., Miranda-Barrientos, J., and Morales, M. (2016). VTA glutamatergic inputs to nucleus accumbens drive aversion by acting on GABAergic interneurons. Nat Neurosci 19, 725–733. 10.1038/nn.4281.

23. Zhang, S., Qi, J., Li, X., Wang, H.L., Britt, J.P., Hoffman, A.F., Bonci, A., Lupica, C.R., and Morales, M. (2015). Dopaminergic and glutamatergic microdomains in a subset of rodent mesoaccumbens axons. Nat Neurosci 18, 386–392. 10.1038/nn.3945.

24. Ernst, M., Zametkin, A.J., Matochik, J.A., Pascualvaca, D., and Cohen, R.M. (1997). Low medial prefrontal dopaminergic activity in autistic children. Lancet 350, 638. 10.1016/s0140-6736(05)63326-0.

25. Nakamura, K., Sekine, Y., Ouchi, Y., Tsujii, M., Yoshikawa, E., Futatsubashi, M., Tsuchiya, K.J., Sugihara, G., Iwata, Y., Suzuki, K., et al. (2010). Brain serotonin and dopamine transporter bindings in adults with high-functioning autism. Archives of general psychiatry 67, 59–68. 10.1001/archgenpsychiatry.2009.137.

26. Zurcher, N.R., Walsh, E.C., Phillips, R.D., Cernasov, P.M., Tseng, C.J., Dharanikota, A., Smith, E., Li, Z., Kinard, J.L., Bizzell, J.C., et al. (2021). A simultaneous [(11)C]raclopride positron emission tomography and functional magnetic resonance imaging investigation of striatal dopamine binding in autism. Transl Psychiatry 11, 33. 10.1038/s41398-020-01170-0.

27. De Rubeis, S., Siper, P.M., Durkin, A., Weissman, J., Muratet, F., Halpern, D., Trelles, M.D.P., Frank, Y., Lozano, R., Wang, A.T., et al. (2018). Delineation of the genetic and clinical spectrum of Phelan-McDermid syndrome caused by SHANK3 point mutations. Molecular autism 9, 31. 10.1186/s13229-018-0205-9.

28. Harony-Nicolas, H., De Rubeis, S., Kolevzon, A., and Buxbaum, J.D. (2015). Phelan McDermid Syndrome: From Genetic Discoveries to Animal Models and Treatment. J Child Neurol. 10.1177/0883073815600872.

29. Boeckers, T.M., Bockmann, J., Kreutz, M.R., and Gundelfinger, E.D. (2002). ProSAP/Shank proteins - a family of higher order organizing molecules of the postsynaptic density with an emerging role in human neurological disease. Journal of neurochemistry 81, 903–910.

30. Filice, F., Vorckel, K.J., Sungur, A.O., Wohr, M., and Schwaller, B. (2016). Reduction in parvalbumin expression not loss of the parvalbumin-expressing GABA interneuron subpopulation in genetic parvalbumin and shank mouse models of autism. Mol Brain 9, 10. 10.1186/s13041-016-0192-8.

31. Gundelfinger, E.D., Boeckers, T.M., Baron, M.K., and Bowie, J.U. (2006). A role for zinc in postsynaptic density asSAMbly and plasticity? Trends in biochemical sciences 31, 366–373.

32. Kreienkamp, H.J. (2008). Scaffolding proteins at the postsynaptic density: shank as the architectural framework. Handbook of experimental pharmacology, 365–380. 10.1007/978-3-540-72843-6_15.

33. Pagano, J., Landi, S., Stefanoni, A., Nardi, G., Albanesi, M., Bauer, H.F., Pracucci, E., Schon, M., Ratto, G.M., Boeckers, T.M., et al. (2023). Shank3 deletion in PV neurons is associated with abnormal behaviors and neuronal functions that are rescued by increasing GABAergic signaling. Molecular autism 14, 28. 10.1186/s13229-023-00557-2.

34. Peca, J., Feliciano, C., Ting, J.T., Wang, W., Wells, M.F., Venkatraman, T.N., Lascola, C.D., Fu, Z., and Feng, G. (2011). Shank3 mutant mice display autistic-like behaviours and striatal dysfunction. Nature 472, 437–442. 10.1038/nature09965.

35. Verpelli, C., Dvoretskova, E., Vicidomini, C., Rossi, F., Chiappalone, M., Schoen, M., Di Stefano, B., Mantegazza, R., Broccoli, V., Bockers, T.M., et al. (2011). Importance of Shank3 protein in regulating metabotropic glutamate receptor 5 (mGluR5) expression and signaling at synapses. The Journal of biological chemistry 286, 34839–34850. 10.1074/jbc.M111.258384.

36. Kolevzon, A., Angarita, B., Bush, L., Wang, A.T., Frank, Y., Yang, A., Rapaport, R., Saland, J., Srivastava, S., Farrell, C., et al. (2014). Phelan-McDermid syndrome: a review of the literature and practice parameters for medical assessment and monitoring. J Neurodev Disord 6, 39. 10.1186/1866-1955-6-39.

37. Levy, T., Foss-Feig, J.H., Betancur, C., Siper, P.M., Trelles-Thorne, M.D.P., Halpern, D., Frank, Y., Lozano, R., Layton, C., Britvan, B., et al. (2022). Strong evidence for genotype-phenotype correlations in Phelan-McDermid syndrome: results from the developmental synaptopathies consortium. Human molecular genetics 31, 625–637. 10.1093/hmg/ddab280.

38. Kolevzon, A., Angarita, B., Bush, L., Wang, A.T., Frank, Y., Yang, A., Rapaport, R., Saland, J., Srivastava, S., Farrell, C., et al. (2014). Phelan-McDermid syndrome: a review of the literature and practice parameters for medical assessment and monitoring. Journa lof Neurodevelopmental Disorders 6.

39. Soorya, L., Kolevzon, A., Zweifach, J., Lim, T., Dobry, Y., Schwartz, L., Frank, Y., Wang, A.T., Cai, G., Parkhomenko, E., et al. (2013). Prospective investigation of autism and genotype-phenotype correlations in 22q13 deletion syndrome and SHANK3 deficiency. Molecular autism 4, 18. 10.1186/2040-2392-4-18.

40. Denayer, A., Van Esch, H., de Ravel, T., Frijns, J.P., Van Buggenhout, G., Vogels, A., Devriendt, K., Geutjens, J., Thiry, P., and Swillen, A. (2012). Neuropsychopathology in 7 Patients with the 22q13 Deletion Syndrome: Presence of Bipolar Disorder and Progressive Loss of Skills. Molecular syndromology 3, 14–20. 10.1159/000339119.

41. Verhoeven, W.M., Egger, J.I., Willemsen, M.H., de Leijer, G.J., and Kleefstra, T. (2012). Phelan-McDermid syndrome in two adult brothers: atypical bipolar disorder as its psychopathological phenotype? Neuropsychiatric disease and treatment 8, 175–179. 10.2147/NDT.S30506.

42. Verhoeven, W.M.A., Egger, J.I.M., and de Leeuw, N. (2020). A longitudinal perspective on the pharmacotherapy of 24 adult patients with Phelan McDermid syndrome. European journal of medical genetics 63, 103751. 10.1016/j.ejmg.2019.103751.

43. Vucurovic, K., Landais, E., Delahaigue, C., Eutrope, J., Schneider, A., Leroy, C., Kabbaj, H., Motte, J., Gaillard, D., Rolland, A.C., and Doco-Fenzy, M. (2012). Bipolar affective disorder and early dementia onset in a male patient with SHANK3 deletion. European journal of medical genetics. 10.1016/j.ejmg.2012.07.009.

44. Leblond, C.S., Nava, C., Polge, A., Gauthier, J., Huguet, G., Lumbroso, S., Giuliano, F., Stordeur, C., Depienne, C., Mouzat, K., et al. (2014). Meta-analysis of SHANK Mutations in Autism Spectrum Disorders: a gradient of severity in cognitive impairments. PLoS genetics 10, e1004580. 10.1371/journal.pgen.1004580.

45. Moessner, R., Marshall, C.R., Sutcliffe, J.S., Skaug, J., Pinto, D., Vincent, J., Zwaigenbaum, L., Fernandez, B., Roberts, W., Szatmari, P., and Scherer, S.W. (2007). Contribution of SHANK3 mutations to autism spectrum disorder. American journal of human genetics 81, 1289–1297. 10.1086/522590.

46. R Development Core Team (2006) R: a language and environment for statistical computing. (2007). R Foundation for Statistical Computing, Vienna, Austria.

47. Bariselli, S., Tzanoulinou, S., Glangetas, C., Prevost-Solie, C., Pucci, L., Viguie, J., Bezzi, P., O’Connor, E.C., Georges, F., Luscher, C., and Bellone, C. (2016). SHANK3 controls maturation of social reward circuits in the VTA. Nat Neurosci 19, 926–934. 10.1038/nn.4319.

48. Bozdagi, O., Sakurai, T., Papapetrou, D., Wang, X., Dickstein, D.L., Takahashi, N., Kajiwara, Y., Yang, M., Katz, A.M., Scattoni, M.L., et al. (2010). Haploinsufficiency of the autism-associated Shank3 gene leads to deficits in synaptic function, social interaction, and social communication. Molecular autism 1, 15. 10.1186/2040-2392-1-15.

49. Harony-Nicolas, H., Kay, M., Hoffmann, J.D., Klein, M.E., Bozdagi-Gunal, O., Riad, M., Daskalakis, N.P., Sonar, S., Castillo, P.E., Hof, P.R., et al. (2017). Oxytocin improves behavioral and electrophysiological deficits in a novel Shank3-deficient rat. Elife 6. 10.7554/eLife.18904.

50. Kouser, M., Speed, H.E., Dewey, C.M., Reimers, J.M., Widman, A.J., Gupta, N., Liu, S., Jaramillo, T.C., Bangash, M., Xiao, B., et al. (2013). Loss of predominant Shank3 isoforms results in hippocampus-dependent impairments in behavior and synaptic transmission. The Journal of neuroscience: the official journal of the Society for Neuroscience 33, 18448–18468. 10.1523/JNEUROSCI.3017-13.2013.

51. Wang, X., McCoy, P.A., Rodriguiz, R.M., Pan, Y., Je, H.S., Roberts, A.C., Kim, C.J., Berrios, J., Colvin, J.S., Bousquet-Moore, D., et al. (2011). Synaptic dysfunction and abnormal behaviors in mice lacking major isoforms of Shank3. Human molecular genetics 20, 3093–3108. 10.1093/hmg/ddr212.

52. Yang, M., Bozdagi, O., Scattoni, M.L., Wohr, M., Roullet, F.I., Katz, A.M., Abrams, D.N., Kalikhman, D., Simon, H., Woldeyohannes, L., et al. (2012). Reduced excitatory neurotransmission and mild autism-relevant phenotypes in adolescent Shank3 null mutant mice. The Journal of neuroscience: the official journal of the Society for Neuroscience 32, 6525–6541. 10.1523/JNEUROSCI.6107-11.2012.

53. Hamdan, F.F., Gauthier, J., Araki, Y., Lin, D.T., Yoshizawa, Y., Higashi, K., Park, A.R., Spiegelman, D., Dobrzeniecka, S., Piton, A., et al. (2011). Excess of de novo deleterious mutations in genes associated with glutamatergic systems in nonsyndromic intellectual disability. American journal of human genetics 88, 306–316. 10.1016/j.ajhg.2011.02.001.

54. Chen, T.W., Wardill, T.J., Sun, Y., Pulver, S.R., Renninger, S.L., Baohan, A., Schreiter, E.R., Kerr, R.A., Orger, M.B., Jayaraman, V., et al. (2013). Ultrasensitive fluorescent proteins for imaging neuronal activity. Nature 499, 295–300. 10.1038/nature12354.

55. Chow, B.Y., Han, X., Dobry, A.S., Qian, X., Chuong, A.S., Li, M., Henninger, M.A., Belfort, G.M., Lin, Y., Monahan, P.E., and Boyden, E.S. (2010). High-performance genetically targetable optical neural silencing by light-driven proton pumps. Nature 463, 98–102. 10.1038/nature08652.

56. Kutlu, M.G., Tat, J., Christensen, B.A., Zachry, J.E., and Calipari, E.S. (2023). Dopamine release at the time of a predicted aversive outcome causally controls the trajectory and expression of conditioned behavior. Cell Rep 42, 112948. 10.1016/j.celrep.2023.112948.

57. Mattis, J., Tye, K.M., Ferenczi, E.A., Ramakrishnan, C., O’Shea, D.J., Prakash, R., Gunaydin, L.A., Hyun, M., Fenno, L.E., Gradinaru, V., et al. (2011). Principles for applying optogenetic tools derived from direct comparative analysis of microbial opsins. Nature methods 9, 159–172. 10.1038/nmeth.1808.

58. Miguel Telega, L., Ashouri Vajari, D., Stieglitz, T., Coenen, V.A., and Dobrossy, M.D. (2022). New Insights into In Vivo Dopamine Physiology and Neurostimulation: A Fiber Photometry Study Highlighting the Impact of Medial Forebrain Bundle Deep Brain Stimulation on the Nucleus Accumbens. Brain Sci 12. 10.3390/brainsci12081105.

59. Parker, K.E., Pedersen, C.E., Gomez, A.M., Spangler, S.M., Walicki, M.C., Feng, S.Y., Stewart, S.L., Otis, J.M., Al-Hasani, R., McCall, J.G., et al. (2019). A Paranigral VTA Nociceptin Circuit that Constrains Motivation for Reward. Cell 178, 653–671 e619. 10.1016/j.cell.2019.06.034.

60. Song, Y., Meng, Q.X., Wu, K., Hua, R., Song, Z.J., Song, Y., Qin, X., Cao, J.L., and Zhang, Y.M. (2020). Disinhibition of PVN-projecting GABAergic neurons in AV region in BNST participates in visceral hypersensitivity in rats. Psychoneuroendocrinology 117, 104690. 10.1016/j.psyneuen.2020.104690.

61. Netser, S., Meyer, A., Magalnik, H., Zylbertal, A., de la Zerda, S.H., Briller, M., Bizer, A., Grinevich, V., and Wagner, S. (2020). Distinct dynamics of social motivation drive differential social behavior in laboratory rat and mouse strains. Nat Commun 11, 5908. 10.1038/s41467-020-19569-0.

62. Hochgerner, H., Singh, S., Tibi, M., Lin, Z., Skarbianskis, N., Admati, I., Ophir, O., Reinhardt, N., Netser, S., Wagner, S., and Zeisel, A. (2023). Neuronal types in the mouse amygdala and their transcriptional response to fear conditioning. Nat Neurosci. 10.1038/s41593-023-01469-3.

63. Netser, S., Haskal, S., Magalnik, H., and Wagner, S. (2017). A novel system for tracking social preference dynamics in mice reveals sex- and strain-specific characteristics. Molecular autism 8, 53. 10.1186/s13229-017-0169-1.

64. Adelsberger, H., Garaschuk, O., and Konnerth, A. (2005). Cortical calcium waves in resting newborn mice. Nat Neurosci 8, 988–990. 10.1038/nn1502.

65. Cui, G., Jun, S.B., Jin, X., Luo, G., Pham, M.D., Lovinger, D.M., Vogel, S.S., and Costa, R.M. (2014). Deep brain optical measurements of cell type-specific neural activity in behaving mice. Nature protocols 9, 1213–1228. 10.1038/nprot.2014.080.

66. Gunaydin, L.A., Grosenick, L., Finkelstein, J.C., Kauvar, I.V., Fenno, L.E., Adhikari, A., Lammel, S., Mirzabekov, J.J., Airan, R.D., Zalocusky, K.A., et al. (2014). Natural neural projection dynamics underlying social behavior. Cell 157, 1535–1551. 10.1016/j.cell.2014.05.017.

67. Molinoff, P.B., and Axelrod, J. (1971). Biochemistry of catecholamines. Annu Rev Biochem 40, 465–500. 10.1146/annurev.bi.40.070171.002341.

68. Solie, C., Girard, B., Righetti, B., Tapparel, M., and Bellone, C. (2022). VTA dopamine neuron activity encodes social interaction and promotes reinforcement learning through social prediction error. Nat Neurosci 25, 86–97. 10.1038/s41593-021-00972-9.

69. Reppucci, C.J., and Veenema, A.H. (2020). The social versus food preference test: A behavioral paradigm for studying competing motivated behaviors in rodents. MethodsX 7, 101119. 10.1016/j.mex.2020.101119.

70. Sun, F., Zeng, J., Jing, M., Zhou, J., Feng, J., Owen, S.F., Luo, Y., Li, F., Wang, H., Yamaguchi, T., et al. (2018). A Genetically Encoded Fluorescent Sensor Enables Rapid and Specific Detection of Dopamine in Flies, Fish, and Mice. Cell 174, 481–496 e419. 10.1016/j.cell.2018.06.042.

71. Srivastava, S., Condy, E., Carmody, E., Filip-Dhima, R., Kapur, K., Bernstein, J.A., Berry-Kravis, E., Powell, C.M., Soorya, L., Thurm, A., et al. (2021). Parent-reported measure of repetitive behavior in Phelan-McDermid syndrome. J Neurodev Disord 13, 53. 10.1186/s11689-021-09398-7.

72. Tavassoli, T., Layton, C., Levy, T., Rowe, M., George-Jones, J., Zweifach, J., Lurie, S., Buxbaum, J.D., Kolevzon, A., and Siper, P.M. (2021). Sensory Reactivity Phenotype in Phelan-McDermid Syndrome Is Distinct from Idiopathic ASD. Genes (Basel) 12. 10.3390/genes12070977.

73. Wang, A.T., Lim, T., Jamison, J., Bush, L., Soorya, L.V., Tavassoli, T., Siper, P.M., Buxbaum, J.D., and Kolevzon, A. (2016). Neural selectivity for communicative auditory signals in Phelan-McDermid syndrome. J Neurodev Disord 8, 5. 10.1186/s11689-016-9138-9.

74. Guillory, S.B., Baskett, V.Z., Grosman, H.E., McLaughlin, C.S., Isenstein, E.L., Wilkinson, E., Weissman, J., Britvan, B., Trelles, M.P., Halpern, D.B., et al. (2021). Social visual attentional engagement and memory in Phelan-McDermid syndrome and autism spectrum disorder: a pilot eye tracking study. J Neurodev Disord 13, 58. 10.1186/s11689-021-09400-2.

75. Richards, C., Powis, L., Moss, J., Stinton, C., Nelson, L., and Oliver, C. (2017). Prospective study of autism phenomenology and the behavioural phenotype of Phelan-McDermid syndrome: comparison to fragile X syndrome, Down syndrome and idiopathic autism spectrum disorder. J Neurodev Disord 9, 37. 10.1186/s11689-017-9217-6.

76. Silverman, J.L., Thurm, A., Ethridge, S.B., Soller, M.M., Petkova, S.P., Abel, T., Bauman, M.D., Brodkin, E.S., Harony-Nicolas, H., Wohr, M., and Halladay, A. (2022). Reconsidering animal models used to study autism spectrum disorder: Current state and optimizing future. Genes, brain, and behavior 21, e12803. 10.1111/gbb.12803.

77. Beier, K. (2022). Modified viral-genetic mapping reveals local and global connectivity relationships of ventral tegmental area dopamine cells. Elife 11. 10.7554/eLife.76886.

78. Beier, K.T., Steinberg, E.E., DeLoach, K.E., Xie, S., Miyamichi, K., Schwarz, L., Gao, X.J., Kremer, E.J., Malenka, R.C., and Luo, L. (2015). Circuit Architecture of VTA Dopamine Neurons Revealed by Systematic Input-Output Mapping. Cell 162, 622–634. 10.1016/j.cell.2015.07.015.

79. Soden, M.E., Chung, A.S., Cuevas, B., Resnick, J.M., Awatramani, R., and Zweifel, L.S. (2020). Anatomic resolution of neurotransmitter-specific projections to the VTA reveals diversity of GABAergic inputs. Nat Neurosci 23, 968–980. 10.1038/s41593-020-0657-z.

80. Bouarab, C., Thompson, B., and Polter, A.M. (2019). VTA GABA Neurons at the Interface of Stress and Reward. Front Neural Circuits 13, 78. 10.3389/fncir.2019.00078.

81. Nieh, E.H., Vander Weele, C.M., Matthews, G.A., Presbrey, K.N., Wichmann, R., Leppla, C.A., Izadmehr, E.M., and Tye, K.M. (2016). Inhibitory Input from the Lateral Hypothalamus to the Ventral Tegmental Area Disinhibits Dopamine Neurons and Promotes Behavioral Activation. Neuron 90, 1286–1298. 10.1016/j.neuron.2016.04.035.

82. Rojek-Sito, K., Meyza, K., Ziegart-Sadowska, K., Nazaruk, K., Puscian, A., Hamed, A., Kielbinski, M., Solecki, W., and Knapska, E. (2023). Optogenetic and chemogenetic approaches reveal differences in neuronal circuits that mediate initiation and maintenance of social interaction. PLoS biology 21, e3002343. 10.1371/journal.pbio.3002343.

83. Ohta, Y., Murakami, T.E., Kawahara, M., Haruta, M., Takehara, H., Tashiro, H., Sasagawa, K., Ohta, J., Akay, M., and Akay, Y.M. (2022). Investigating the Influence of GABA Neurons on Dopamine Neurons in the Ventral Tegmental Area Using Optogenetic Techniques. Int J Mol Sci 23. 10.3390/ijms23031114.

84. Robinson, D.L., Heien, M.L., and Wightman, R.M. (2002). Frequency of dopamine concentration transients increases in dorsal and ventral striatum of male rats during introduction of conspecifics. J Neurosci 22, 10477–10486.

85. Robinson, D.L., Zitzman, D.L., Smith, K.J., and Spear, L.P. (2011). Fast dopamine release events in the nucleus accumbens of early adolescent rats. Neuroscience 176, 296–307. 10.1016/j.neuroscience.2010.12.016.

